# Positive allosteric modulation of emodepside sensitive *Brugia malayi* SLO-1F and *Onchocerca volvulus* SLO-1A potassium channels by GoSlo-SR-5-69

**DOI:** 10.1101/2025.01.31.635861

**Authors:** Mark McHugh, Charity N Njeshi, Nathaniel Smith, Sudhanva S. Kashyap, Real Datta, Han Sun, Alan P. Robertson, Richard J. Martin

**Affiliations:** Department of Biomedical Sciences, College of Veterinary Medicine, Iowa State University, Ames, IA, USA; Leibniz-Forschungsinstitut fur Molekulare Pharmacologie (FMP), Berlin, Germany; Institute of Chemistry, Technical University of Berlin, Germany

**Keywords:** Emodepside, GoSlo-SR-5-69, lymphatic filariasis, onchocerciasis, *Brugia malayi*, *Onchocerca volvulus*, SLO-1 K

## Abstract

Human lymphatic filariasis and onchocerciasis are Neglected Tropical Diseases (NTDs), of major public health concern. Prophylaxis and treatment rely on anthelmintics that effectively eliminate migrating microfilariae but lack efficacy against adult filarial worms. To expedite the elimination of both diseases, the introduction of drugs with adulticidal activity is paramount. The broad-spectrum anthelmintic emodepside, a nematode selective SLO-1 K channel activator, has been considered a promising candidate for the treatment of onchocerciasis due to its macrofilaricidal activity against *Onchocerca volvulus*. Nevertheless, it is less effective against adult *Brugia malayi,* one of the causative agents of human lymphatic filariasis. Characterizing molecular and pharmacological disparities between highly conserved splice variant isoforms of *B. malayi* and *O. volvulus* SLO-1 K channels and identifying allosteric modulators that can increase emodepside potency on *B. malayi* SLO-1 K channels is necessary for therapeutic advance. In this study, we tested the effects of emodepside and the mammalian BK channel activator, GoSlo-SR-5-69 alone and in combination on *Xenopus* expressed *B. malayi* SLO-1F and *O. volvulus* SLO-1A channels. Additionally, binding poses of emodepside, and GoSlo-SR-5-69 were predicted on both channels using molecular docking. *Ovo*-SLO-1A was more sensitive to emodepside than *Bma*-SLO-1F. GoSlo-SR- 5-69 was a positive allosteric modulator, potentiating the effects of emodepside. Emodepside was docked at the S6 pocket below the selectivity filter for *Bma*-SLO-1F and *Ovo*-SLO-1A. The binding of emodepside in the S6 pocket indicated a stabilizing *π-π* interaction between F342 and the phenyl rings of emodepside which may contribute to the potency of emodepside on these filaria channels. Molecular docking suggested that GoSlo-SR-5-69 binds at the RCK1 pocket. This study reveals for the first time, allosteric modulation of filarial nematode SLO-1 K channels by a mammalian BK channel activator and highlights its ability to increase emodepside potency on the *B. malayi* SLO-1 K channel.

**Author Summary:** *B. malayi* is one of the causative agents of lymphatic filariasis, while *O. volvulus* causes onchocerciasis. Elimination of adult *B. malayi* and *O. volvulus* is impeded by anthelmintics lacking adequate macrofilaricidal activity. The anthelmintic emodepside has microfilaricidal and macrofilaricidal activity. However, it exhibits poor efficacy against adult *B. malayi in vivo.* Here, we conducted a comparative pharmacological characterization of two closely related splice variants SLO-1 K channels of *B. malayi* and *O. volvulus*. Using voltage clamp, we tested the effects of emodepside and GoSlo-SR-5-69 independently and in combination on *Xenopus* expressed *B. malayi* SLO-1F and *O. volvulus* SLO-1A channels. Binding poses of emodepside, and GoSlo-SR-5-69 were predicted on both channels using molecular docking. We demonstrate that the *O. volvulus* SLO-1A receptor is more sensitive to emodepside than the *B. malayi* SLO-1F receptor. GoSlo-SR-5-69 is also a potentiator of emodepside responses. Homology modelling revealed that emodepside probably binds in the pore region of the B. *malayi* SLO-1F and *O. volvulus* SLO-1A channels, while GoSlo-SR-5-69 probably binds to the RCK1 domain of the channels. Our study provides new insights into similarities and differences shared between SLO-1 K channels of each species and reveals positive allosteric modulation by a mammalian SLO-1 K activator.

## Introduction

Onchocerciasis (river blindness) and lymphatic filariasis (elephantiasis), are diseases that have significantly impaired the health of individuals in tropical regions. Lymphatic filariasis is caused by 3 filarial parasites, namely, *Brugia malayi*, *Wuchereria bancrofti* and *Brugia timori,* while onchocerciasis is caused by the nematode *Onchocerca volvulus* [1]. Approximately 51 million people are reported to be infected with lymphatic filariasis, while 20.9 million individuals are infected with onchocerciasis [2].

Mass drug administration (MDA) programs have been implemented for prevention and treatment of these infections by use of anthelmintics such as ivermectin [3], albendazole [4] and diethylcarbamazine [5]. These drugs have effects on infectious stages of the parasites, but lack sufficient macrofilaricidal activity, thus spurring the need for the development of effective drugs. The semi-synthetic cyclooctadepsipeptide, emodepside, has shown inhibitory effects on various developmental stages of filarial nematodes of human and veterinary importance [6, 7]. Its mode of action involves the selective activation of the calcium-activated K^+^ channel (SLO-1 K) of nematodes, leading to inhibition of locomotion and complete paralysis [8–10]. Therefore, emodepside is a valuable ingredient for therapeutic use in human medicine and has completed phase II clinical trials [11–13].

Despite the development of effective drugs, decreased efficacy generally occurs with wide-spread use, due to the emergence of drug resistance. Hence, delaying the onset of resistance is paramount and may be achieved by a combination of two or more drugs that have additive or synergistic effects. Potential candidates for combination with emodepside are the mammalian BK channel openers. One example of the BK channel openers that we found to be active on our filarial SLO-1 K channels was GoSlo-SR-5-69. This compound belongs to the GoSlo-SR family of compounds that have been studied for their effects on overactive bladder dysfunction in experimental animals [14].

We compared pharmacological differences between the structurally related splice variant channels of *Brugia malayi* (SLO-1F) and *Onchocerca volvulus* (SLO-1A) and their interaction with emodepside and GoSlo-SR-5-69 alone and in combination. Emodepside activated both channels in a concentration-dependent manner with *Ovo*-SLO-1A showing higher sensitivity to emodepside than *Bma*-SLO-1F. Additionally, emodepside reduced the voltage-activation of both channels. 3 µM GoSlo-SR-5-69 alone did not activate either of the channels, but acted as a positive allosteric modulator, potentiating 0.3 µM emodepside. Our molecular docking studies suggest that emodepside binds in the S6 pocket, just below the selectivity filter for emodepside in *Bma*-SLO-1F and *Ovo*-SLO-1A, while GoSlo-SR-5-69 is suggested to bind in the RCK1 pocket A which appears only in he open state of the *Drosophila melanogaster* SLO-1 K channel [15]. These findings highlight similarities and differences between the two filarial nematode SLO-1K channels and suggest a proof-of-concept approach for increasing emodepside potency.

## Results

### *Bma*-SLO-1F and *Ovo*-SLO-1A are highly conserved

The *Brugia malayi* genome has two *slo-1* isoforms (*slo-1a* and *slo-1f*), whereas the *Onchocerca volvulus* genome has five isoforms (*slo-1a, b, c, d,* and *f*). Furthermore, functional expression of these splice variants in the *Xenopus* oocyte expression system have demonstrated pharmacological differences in the sensitivity of *B. malayi slo-1* splice variants to emodepside [7], but the sensitivities to emodepside of the different splice variants of *O. volvulus slo-1* were not reported to be different [16]. We compared the amino acid sequences of *B. malayi* and *O. volvulus slo-1* splice variants, to identify the highest sequence conservation between isoforms of the two filarial parasites. We identified the *B. malayi* SLO-1F and *O. Volvulus* SLO-1A splice variants.

Fig. 1 shows the amino acid sequence alignment of *Bma*-SLO-1F and *Ovo*-SLO- 1A using the EMBOSS Needle pairwise sequence online alignment tool. Annotated are characteristic BK channel domains, namely, the N-terminal transmembrane domain, consisting of a voltage sensor domain (enclosed in orange box) and the pore domain (enclosed in blue box). Additionally, within the pore domain is the selectivity filter (light blue box). Fig 1 also shows the cytosolic tail domain (CTD) that includes the two regulating domains for potassium (K^+^) conductance, RCK1 (pink box) and RCK2 (black box). Overall, the isoforms showed 96.2% identity between the two sequences.

**Fig 1.**
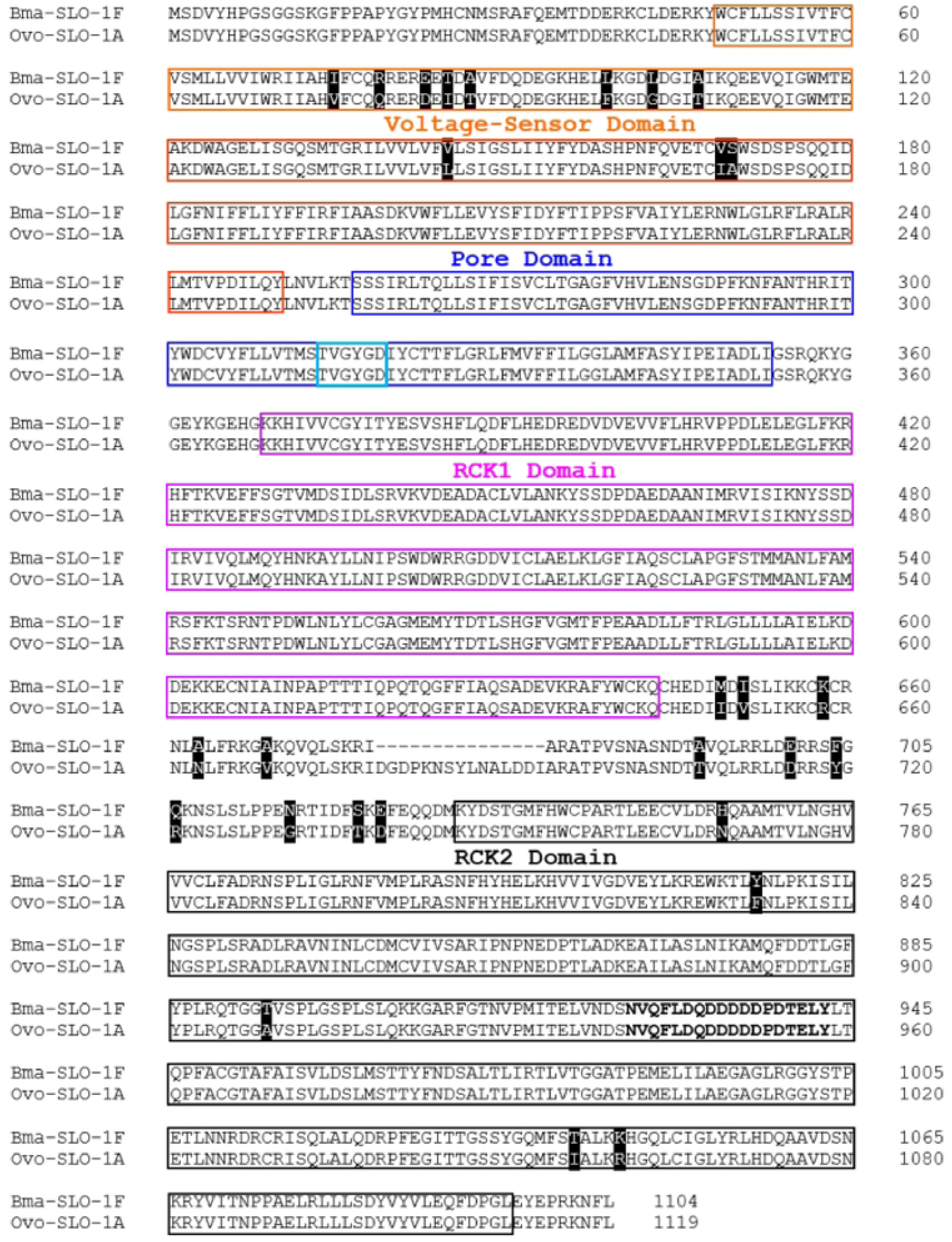
Amino acid sequence alignment of *Bma*-SLO-1F and *Ovo*-SLO-1A. The voltage sensor domain (VSD; orange boxes), pore gate domain (PD; blue boxes) comprising the selectivity filter (light blue box) and two C-terminal domains for regulator of K^+^ conductance (RCK1; pink boxes and RCK2; black boxes) are indicated. Amino acids which are not identical between filarial species are highlighted by a black background. Gaps are indicated by “_” symbols for amino acid residues that are missing.

Further alignment of each individual domain revealed that the voltage sensor domain was 94.5% identical between both isoforms, with differences in eleven (11) amino acid residues (highlighted in black). In contrast, the pore domain and RCK1 domain had identical amino acid residues (100% identity), whereas the RCK2 domain was 98.6% identical with 5 differences in amino acid residues (highlighted in black). The region between the RCK1 and RCK2 domain was 73.5% identical, with *Ovo*-SLO-1A having 15 additional amino acid residues when compared to *Bma*-SLO-1F. There were also 12 residues that were not identical (highlighted in black).

Taken together, both *Bma*-SLO-1F and *Ovo*-SLO-1A are highly conserved in all domains. Nevertheless, of the four domains, the voltage sensor domain is least conserved with 11 amino acid differences which may be critical in influencing the opening and pharmacology of the individual receptors [17]. Furthermore, the presence or absence of residues between the RCK1 and RCK2 domain of *Ovo*-SLO-1A and *Bma*-SLO-1F respectively, may also result in small differences in the receptor conformation during the binding of a ligand or divalent cations and intracellular signaling molecules.

### *Ovo*-SLO-1A is more sensitive to emodepside than *Bma*-SLO-1F

To test for functional expression of our cloned *Bma-slo-1f* and *Ovo-slo-1a* genes in the *Xenopus laevis* oocyte expression system, we conducted cumulative concentration-response experiments as described in the methods. Fig 2A shows representative traces produced from our recordings for *Ovo*-SLO-1A (top; pink trace) and *Bma*-SLO-1F (lower; blue trace) splice variants. We observed that perfusion of increasing concentrations of emodepside (0.1, 0.3, 1, 3, 10 µM), elicited channel activation, with a concentration-dependent increase in outward currents in oocytes expressing their respective ion-channel receptors.

**Fig 2.**
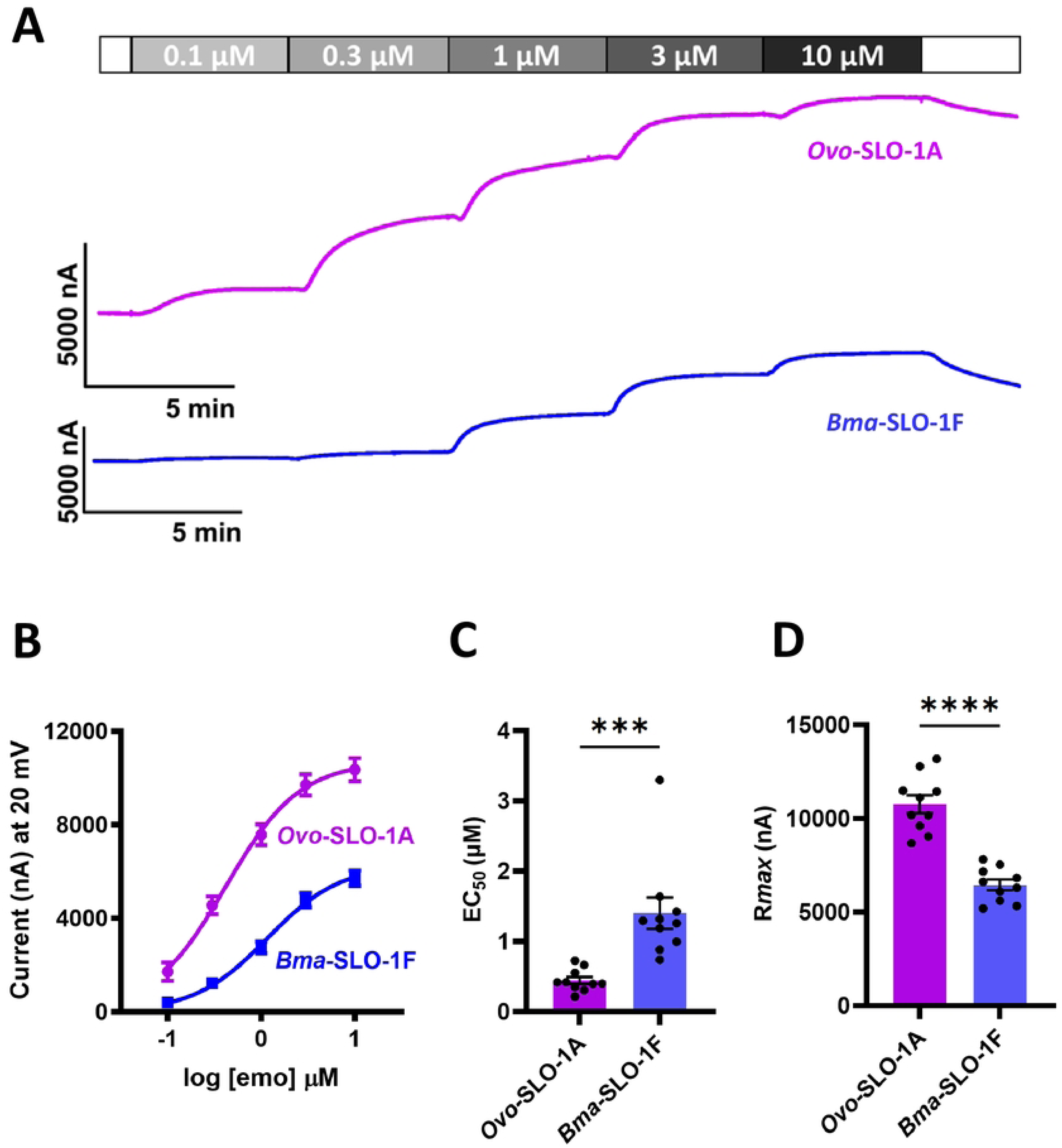
Emodepside (emo) concentration response relationships for *Ovo*-SLO-1A and *Bma*-SLO-1F. **A.** Representative traces for two-electrode voltage-clamp recording showing outward currents for *Ovo*-SLO-1A (top; pink trace) and *Bma*-SLO-1F (lower; blue trace) channels, elicited in the presence of increasing concentrations of emodepside (0.1 to 10 µM) at a holding potential of +20 mV. **B.** Emodepside concentration-response curves for *Ovo*-SLO-1A (n = 10; pink) and *Bma*-SLO-1F (n = 10, blue). **C.** Bar chart (mean ± S.E.M) of the EC_50_ of emodepside for *Ovo*-SLO-1A (n = 10; pink) and *Bma*-SLO-1F (n = 10; blue) channels. The *Ovo*-SLO-1A channel is more sensitive to emodepside than *Bma*-SLO-1F **D:** Bar chart (mean ± S.E.M) of the maximum current responses (*Rmax*) of emodepside for *Ovo*-SLO-1A (n = 10; pink) and *Bma*-SLO-1F (n = 10, blue) channels. Bottom was constrained to zero for curve fitting. The *Ovo*-SLO-1A channel has a higher efficacy than the *Bma*-SLO-1F channel. \*\*\**P* < 0.001, \*\*\*\**P* < 0.0001 significantly different as indicated; unpaired two-tailed student t- test.

Fig 2B shows emodepside concentration-response relationships for both channels. The EC_50_ and maximum response (*R_max_*) values for emodepside for *Ovo*-SLO- 1A expressing oocytes were 0.40 ± 0.05 µM and 10762 ± 478 nA, (n = 10), while the EC_50_ and maximum response (*R_max_*) values for *Bma*-SLO-1F were 1.4 ± 0.2 µM and 6446 ± 287 nA, (n = 10). The EC_50_ for *Ovo*-SLO-1A was significantly smaller than that for *Bma*- SLO-1F, thus suggesting that *Ovo*-SLO-1A is 3.5 times more sensitive to emodepside than *Bma*-SLO-1F (Fig. 2C). Our analysis of the R_max_ values also show that *Ovo*-SLO-1A had a statistically significantly higher value in comparison to *Bma*-SLO-1F (Fig. 2D). Finally, we observed that the Hillslope values for the *Ovo*-SLO-1A expressing oocytes was 1.2 ± 0.1 and oocytes expressing *Bma*-SLO-1F was 1.1 ± 0.1. These values showed little cooperativity suggesting only 1 molecule of emodepside was binding with the SLO- 1 K channels.

### Effects of emodepside on *Ovo*-SLO-1A and *Bma*-SLO-1F current-voltage relationships

To investigate the effects of emodepside on voltage-dependent currents, we conducted voltage step experiments on oocytes expressing *Ovo*-SLO-1A, *Bma*-SLO-1F and water injected oocytes (control). Mean current response for water injected oocytes in the absence or presence of emodepside were similar over the range of step potentials (Fig 3A and C). This showed that emodepside had little or no agonist effects on any endogenous channels of *Xenopus laevis*.

**Fig 3.**
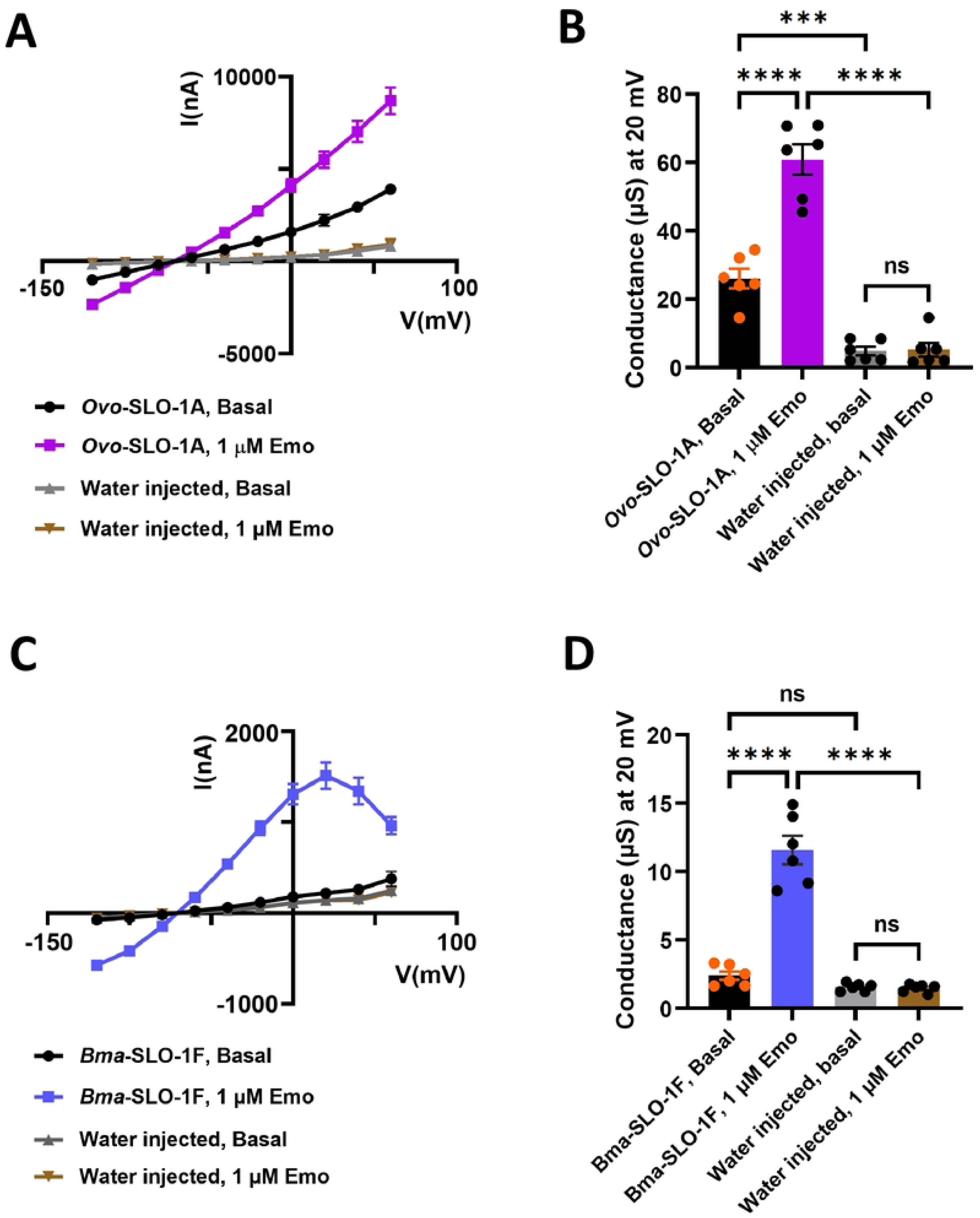
Effects of emodepside on current-voltage curves (IVCs) of the *Ovo*-SLO-1A and *Bma*-SLO-1F channels expressed in *Xenopus laevis* oocytes. **A.** Basal currents (mean ± S.E.M) from oocytes expressing *Ovo*-SLO-1A (n = 6, black) or injected with water (n = 6, grey) in the absence of emodepside. Currents (mean ± S.E.M) obtained from oocytes expressing *Ovo*-SLO-1A (n = 6, pink) or injected with water (n = 6, red) in the presence of 1 µM emodepside. **B.** Bar chart (mean ± S.E.M) of the slope conductance of *Ovo*-SLO-1A expressing oocytes perfused with recording solution and no drug (n = 6; black), *Ovo*-SLO-1A expressing oocytes exposed to 1 µM emo (n = 6; pink), water injected oocytes perfused with recording solution and no drug (grey; n = 6) and water injected oocytes exposed to 1 µM emodepside (red; n = 6). **C.** Basal currents (mean ± S.E.M) from oocytes expressing *Bma*-SLO-1F (n = 6, black) or injected with water (n = 6, grey) in the absence of emodepside. Currents (mean ± S.E.M) obtained from oocytes expressing *Bma*-SLO-1F (n = 6, blue) or injected with water (n = 6, tan) in the presence of 1 µM emodepside. **D.** Bar chart (mean ± S.E.M) of the slope conductance of *Bma*-SLO-1F expressing oocytes perfused with recording solution and no drug (n = 6; black), *Bma*-SLO-1F expressing oocytes exposed to 1 µM emo (n = 6; pink), water injected oocytes perfused with recording solution and no drug (grey; n = 6) and water injected oocytes exposed to 1 µM emodepside (red; n = 6).

However, in the absence of 1 µM emodepside, the slope of current-voltage relationships for oocytes expressing *Ovo*-SLO-1A channels showed an increase in conductance to 26 ± 2.9 µS compared to the water injected controls, 4.81 ± 1.2 µS, with a reversal potential of −71 mV, suggesting that some of the expressed *Ovo*-SLO-1A channels are open at rest (Fig 3A and B). The application of 1 µM emodepside produced an increase in inward current responses at potentials more negative than the reversal potential, −71 mV, and an increase in outward current at potentials positive than the reversal potential, (Fig 3A and B). There was an increase in the conductance to 61 ± 4.4 µS, but no change in the reversal potential (Fig 3A and B). Collectively, these observations demonstrate that emodepside increases the opening of the filarial *Ovo*-SLO- 1A channels expressed in *Xenopus* oocytes.

Unlike the *Ovo-*SLO-1A channels that produced high current responses in the absence and presence of emodepside, *Bma*-SLO-1F channels showed smaller current responses. Firstly, we observed that the control (basal) currents for oocytes expressing *Bma*-SLO-1F were not significantly different from water injected oocytes in the absence of 1 µM emodepside (Fig 3C). The difference in conductance, 2.4 ± 0.3 µS, of the *Bma*- SLO-1F injected oocytes compared to the conductance, 1.6 ± 0.1 µS, of the water injected oocytes was not significant (Fig 3D). This indicates that there were fewer open *Bma*-SLO- 1F channels present in the oocytes.

Secondly, the conductance change induced by the application of 1 µM emodepside was smaller. 1 µM emodepside, increased the conductance from 2.4 ± 0.3 µS to 12 ± 1.0 µS in the *Bma*-SLO-1F expressing oocytes. Again, 1 µM emodepside produced no detectable change in the conductance for water injected oocytes (Fig 3C and D). The reversal potential of all the current voltage plots was close to −72 mV and close to the reversal potential of the *Ovo*-SLO-1A expressed channels.

Thirdly, a reduction in the outward current response was notably different at +40 mV to +60 mV step potentials for *Bma*-SLO-1F (Fig 3C). This phenomenon was absent from the current-voltage curves of *Ovo*-SLO-1A. Despite having very similar sequences, *Ovo*-SLO-1A and *Bma*-SLO-1F, showed clear detectable differences in their current-voltage relationship, suggesting that the differences in the voltage-sensitive regions of the channels affects the response to emodepside.

### Verruculogen inhibits emodepside-induced responses of *Ovo*-SLO-1A and *Bma*- SLO-1F channels

To validate that the observed currents produced by the oocytes expressing *Ovo*- SLO-1A and *Bma*-SLO-1F were due to emodepside activation, the mammalian BK channel blocker, verruculogen was used as an antagonist. We assessed the effects of verruculogen before and after the addition of emodepside to determine whether the order of application influenced the outcome of its antagonist properties.

We observed that application of 1 µM verruculogen to the oocytes alone, significantly reduced the *Ovo*-SLO-1A currents, while having no effects on the voltage dependent currents of *Bma*-SLO-1F (Fig 4A and B). This confirmed our earlier findings that some of the expressed *Ovo*-SLO-1A channels are open at rest, thus producing large basal currents in the absence of the agonist, emodepside. Subsequent application of 1 µM emodepside did not produce channel activation as seen when emodepside was first applied in our previous experiments for both channels (Fig 4A and B). In fact, the currents were like basal currents prior to drug application (Fig. 4A and B).

**Fig 4.**
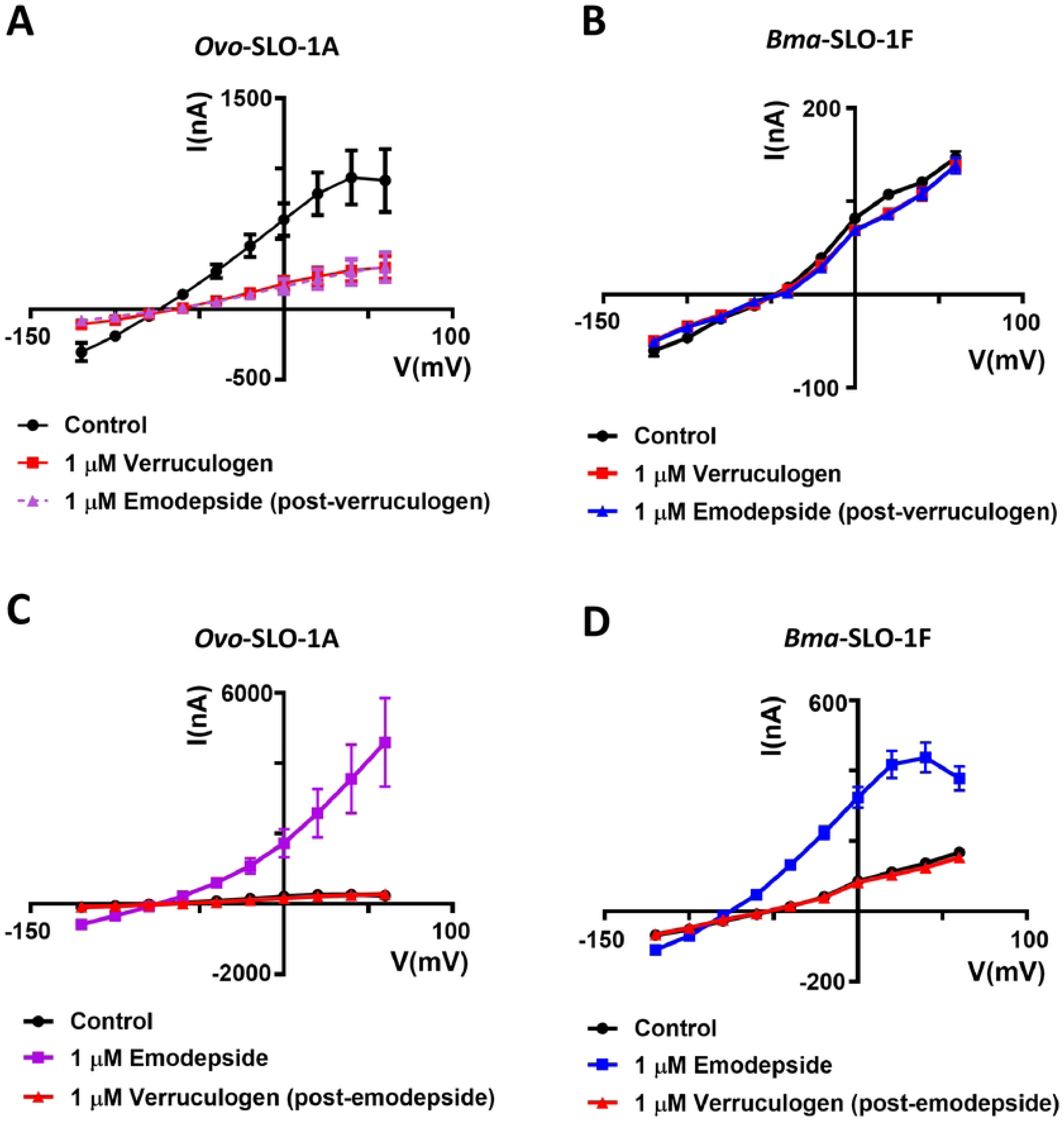
Inhibitory effects of verruculogen on *Ovo*-SLO-1A, *Bma*-SLO-1F and emodepside mediated currents. **A.** Current voltage curves (IVCs) (mean ± S.E.M) corresponding to oocytes expressing the *Ov*o-SLO-1A channel (n = 6), without the addition of drugs (control; indicated as black IVC), after application of 1 µM verruculogen (red IVC) and after application of 1 µM emodepside (magenta IVC). **B.** Currents obtained from oocytes expressing the *Bma*-SLO-1F channel (n = 6) in the absence of drugs (control) indicated in black, in the presence of 1 µM verruculogen indicated in red and subsequently treated with 1 µM emodepside indicated in blue. **C.** Currents obtained from oocytes expressing *Ovo*-SLO-1A (n = 6) in the absence of drugs (control, black) followed by immediate application of 1 µM emodepside (magenta) and subsequently 1 µM verruculogen (red). **D.** Oocytes expressing the *Bma*-SLO-1F channel perfused in the absence of drugs (control, black) or with 1 µM emodepside (blue) or with 1 µM verruculogen (red). For panels **A** and **B**, oocytes were perfused with a recording solution for 1 min before the IVC was recorded. After perfusion with the recording solution, verruculogen was applied for 2 min followed by the recording of IVC. Finally, emodepside was added for 5 min followed by IVC recordings. Panels B and C had a very similar protocol with the exception that emodepside was being added before verruculogen.

Given that verruculogen blocked the activation of the filarial nematode splice variant channels when applied before emodepside, we then determined its effects after emodepside application. As expected, 1 µM emodepside produced activation of both splice variant channels in a similar manner as previously reported in this study. Briefly, emodepside elicited a dramatic increase in currents for *Ovo*-SLO-1A and *Bma*-SLO-1F expressing oocytes (Fig 4C and D). Next, the application of 1 µM verruculogen immediately after emodepside treatment led to a significant decrease in emodepside induced currents (Fig. 4C and D). These results were notable since previous work suggested that the addition of verruculogen after emodepside application does not inhibit emodepside induced currents for the *Cel*-SLO-1A channel [18]. Overall, these findings highlight that in addition to mammalian and *Caenorhabditis elegans* SLO-1 K channels, verruculogen is also an antagonist of filarial nematode SLO-1 K channels.

### GoSlo-SR-5-69 is not an activator of *Bma*-SLO-1F and *Ovo*-SLO-1A channel

To explore the effects of additional synthetic BK channel activators on filarial SLO- 1 K channels, we selected and tested the tetrahydro-2-napthalene derivative, GoSlo-SR-5-69 which is a mammalian BK channel opener. We observed in oocytes held at a steady-state potential of +20 mV, that application of 3 µM GoSlo-SR-5-69 to water injected oocytes and oocytes expressing *Bma*-SLO-1F and *Ovo*-SLO-1A receptors, produced slowly inactivating inward current, (Fig 5A, 5B and 5C). In contrast, subsequent application of 0.3 µM emodepside resulted in the activation of *Bma*-SLO-1F and *Ovo*- SLO-1A channels that produced outward K^+^ currents, (Fig 5B and C). Water injected oocytes yielded no response to emodepside. These results show that GoSlo-SR-5-69 does not by itself activate *Bma*-SLO-1F or *Ovo*-SLO-1A receptors.

**Fig 5.**
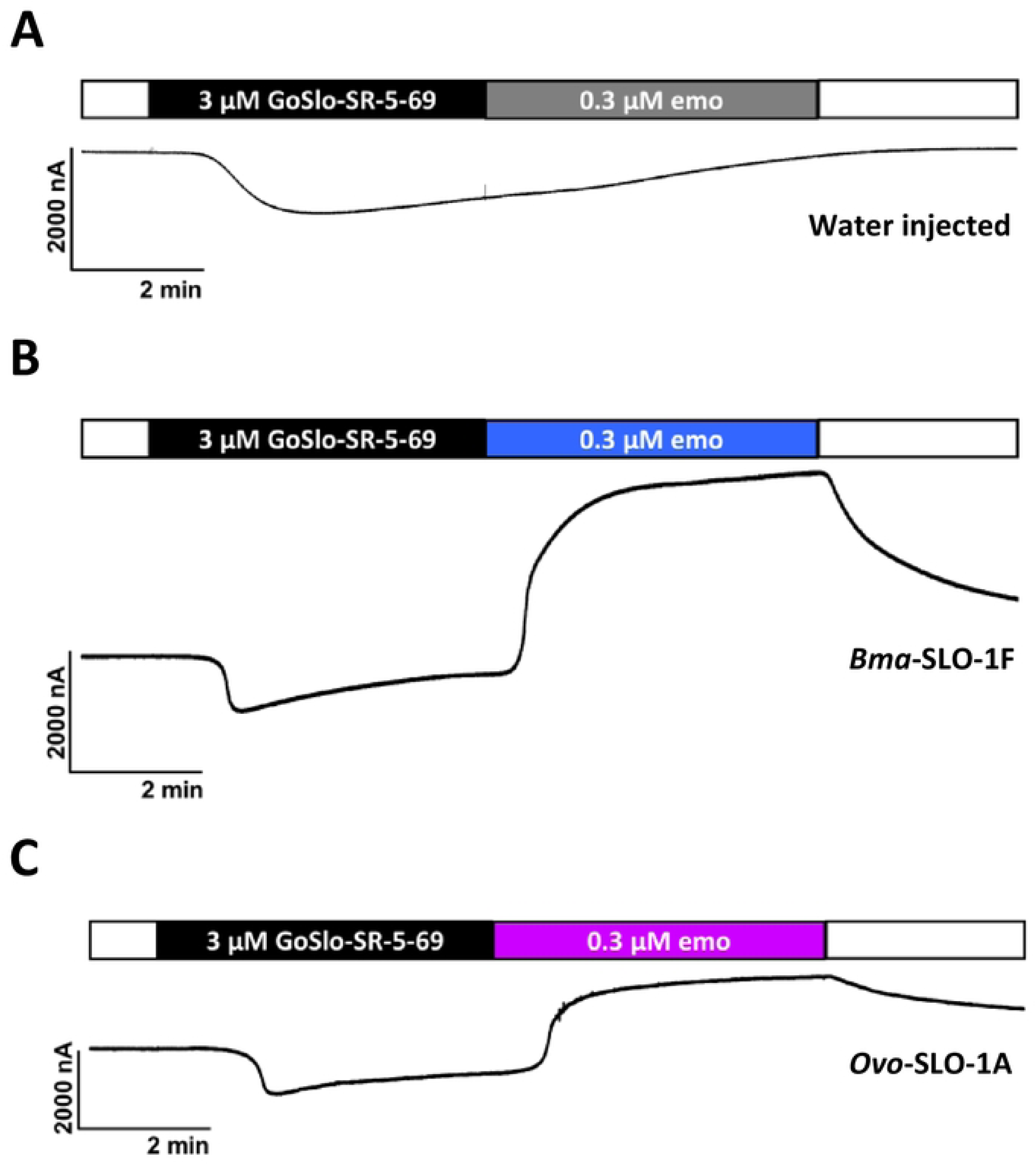
Effects of GoSlo-SR-5-69 on *Bma*-SLO-1F and *Ovo*-SLO-1A channels. **A.** Sample trace of water injected oocytes (n = 6). **B.** Sample trace of oocytes expressing the *Bma*-SLO-1F channel (n = 6). **C.** Sample trace of *Ovo*-SLO-1A expressing channel (n = 6). Application of 3 µM GoSlo-SR-5-69 produced transient inactivating inward currents in oocytes injected with water, *Bma*-SLO-1F and *Ovo*-SLO-1A channels. Emodepside failed to activate water injected oocytes but produced outward currents in oocytes expressing *Bma*-SLO-1F and *Ovo*-SLO-1A channels.

### GoSlo-SR-5-69 is a Positive Allosteric Modulator of filarial nematode SLO-1 channels

Our experiments above demonstrated that GoSlo-SR-5-69 lacks agonist activity on *Bma*-SLO-1F and *Ovo*-SLO-1A receptors at a concentration of 3 µM. To test for any allosteric emodepside modulating effects of GoSlo-SR-5-69, oocytes were perfused with 0.3 µM emodepside for channel activation, followed by co-application of 3 µM GoSlo-SR- 5-69 in the continued presence of emodepside (Fig 6A). This was then followed by washing of the GoSlo-SR-5-69 and then washing of the 0.3 µM emodepside with recording solution.

**Fig 6.**
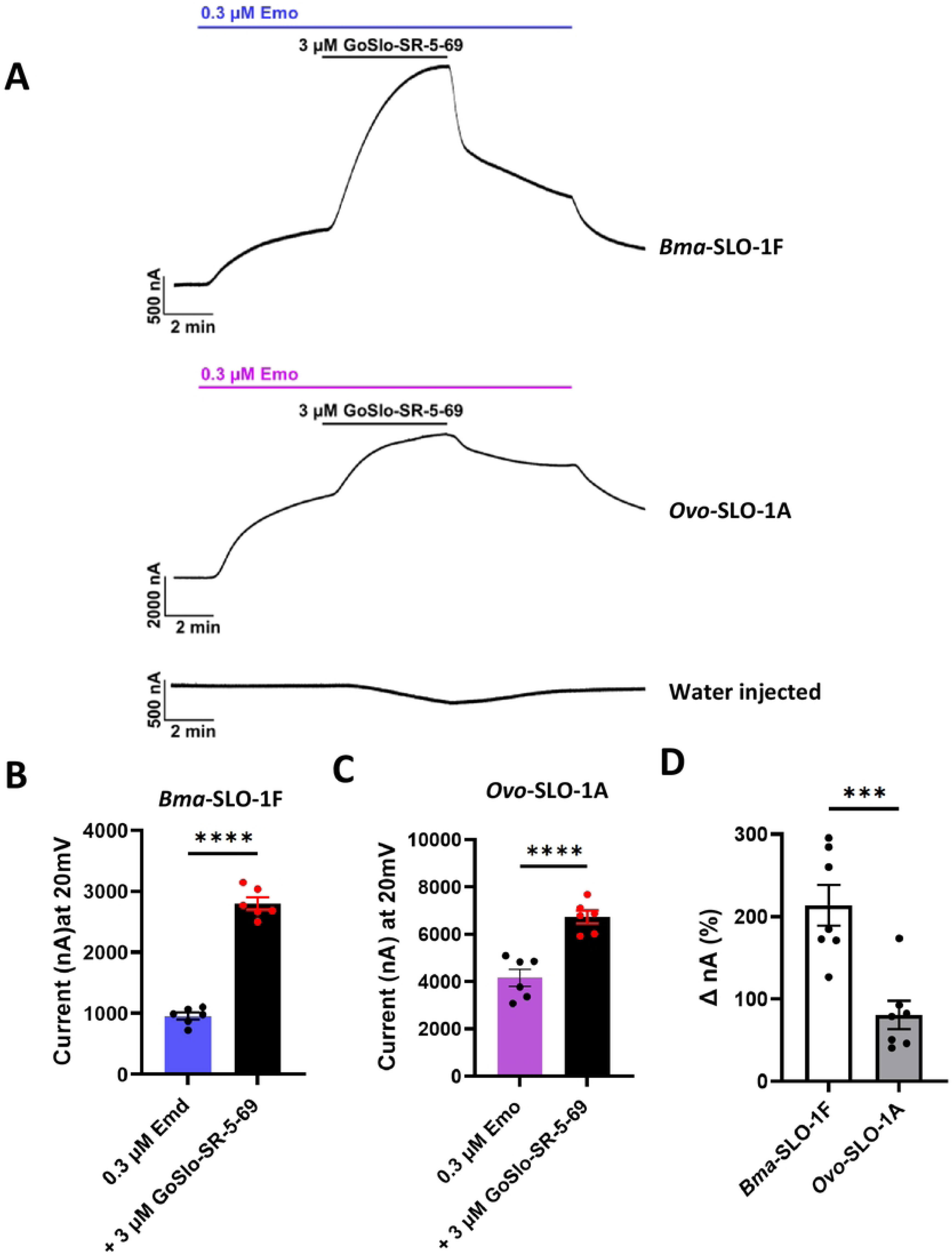
Effects of GoSlo-SR-5-69 on *Bma*-SLO-1F and *Ovo*-SLO-1A-mediated emodepside responses. **A.** Representative traces for *Bma*-SLO-1F, *Ovo*-SLO-1A and water injected oocytes perfused with 0.3 µM emodepside, followed by 3 µM GoSlo-SR-5- 69 in the continued presence of emodepside and an initial wash with 0.3 µM emodepside with a final wash with oocyte recording solution. **B.** Bar chart showing current (in nA) generated in response to 0.3 µM emodepside alone and in combination with 3 µM GoSlo-SR-5-69 for oocytes expressing the *Bma*-SLO-1F channel. Blue bar: 0.3 µM emodepside alone (n = 6). Black bar: 3 µM GoSlo-SR-5-69 co-applied with 0.3 µM emodepside (n = 6). **C.** Bar chart showing current (in nA) generated in response to 0.3 µM emodepside alone and in combination with 3 µM GoSlo-SR-5-69 for oocytes expressing the *Ovo*-SLO- 1A channel. Pink bar: 0.3 µM emodepside alone (n = 6). Black bar: 3 µM GoSlo-SR-5-69 in combination with 0.3 µM emodepside (n = 6). **D.** Bar chart of percentage (%) increase in currents produced by emodepside and GoSlo-SR-5-69 co-application for *Bma*-SLO-1F (white, n = 6) and *Ovo*-SLO-1A (grey, n = 6). Data are plotted as mean ± S.E.M; \*\*\*\**P* < 0.0001 significantly different as indicated; paired two-tailed student t-test; \*\**P* < 0.001, significantly different as indicated, unpaired two-tailed student t-test.

We observed that activating both *Bma*-SLO-1F and *Ovo*-SLO-1A with emodepside (0.3 µM), then co-applying 3 µM GoSlo-SR-5-69 with emodepside, produced an additional increase in the outward current amplitude response for both channels (Fig 6A). Washing off the GoSlo-SR-5-69 reduced the amplitude of the current response of both receptors. Water injected oocytes did not show activation with emodepside or potentiation when GoSlo-SR-5-69 and emodepside were co-applied (Fig 6A). These results provide validation that the observed potentiation of channel openings of *Bma*-SLO-1F and *Ovo*- SLO-1A is elicited by GoSlo-SR-5-69.

Using traces of experiments like those shown in Fig 6A, we quantified current responses for both receptors in the presence of 0.3 µM emodepside alone, and in combination with 3 µM GoSlo-SR-5-69. The current (I_max_) responses for oocytes expressing *Bma*-SLO-1F challenged with 0.3 µM emodepside alone were 954 ± 355 nA, (n = 6); in the combined presence of 3 µM GoSlo-SR-5-69 and 0.3 µM emodepside, they were 2799 ± 100 nA, (n = 6), (Fig 6B). For *Ovo*-SLO-1A injected oocytes, the current responses were 4170 ± 355 nA, (n = 6) in the sole presence of emodepside and 6735 ± 272 nA, (n = 6) in the presence of 3 µM GoSlo-SR-6-69 (Fig 6C). Our analysis confirmed that GoSlo-SR-5-69 significantly increased the emodepside responses of both *Bma*-SLO- 1F and *Ovo*-SLO-1A receptors.

We also calculated the percentage increases in peak currents produced by 3 µM GoSlo-SR-5-69 on the 0.3 µM emodepside response for *Bma*-SLO-1F which was 2.6 times larger than that of *Ovo*-SLO-1A, namely 200% and 65.2% respectively (Fig 6D). These percent increases imply a greater potency of GoSlo-SR-5-69 on *Bma*-SLO-1F. These findings are notable and highlight the positive allosteric effect of GoSlo-SR-5-69 on the action of the anthelmintic emodepside on the SLO-1 K channels of two filarial nematode species that cause ‘river blindness’ and elephantiasis.

### Extracellular Ca^2+^ is not required for GoSlo-SR-5-69 potentiation of emodepside

We have observed that GoSlo-SR-5-69 produces a slowly inactivating inward current in water injected oocytes and oocytes expressing *Bma*-SLO-1F or *Ovo*-SLO-1A channels. In addition, we have also observed that the activation of the filarial nematodes SLO-1 K splice variant channels with emodepside, followed by co-application with GoSlo-SR-5-69 led to a significant potentiation of emodepside responses. If the transient inward currents produced by GoSlo-SR-5-69 were due to activation of Ca^2+^ permeable channels, then the entry of Ca^2+^ may lead to the activation of *Bma*-SLO-1F and *Ovo*-SLO-1A channels. To test this possibility, we replaced Ca^2+^ with equimolar Co^2+^ in our oocyte recording solution to inhibit the entry of Ca^2+^ during our recordings.

The inward current response to GoSlo-SR-5-69 persisted unchanged when extracellular Ca^2+^ was replaced by Co^2+^ in the water injected oocytes and oocytes expressing the *Bma*-SLO-1F or *Ovo*-SLO-1A channels, as shown in Figs S1A,B&C. These currents were not Ca^2+^ currents and emodepside still activated outward currents when Ca^2+^ was replaced by Co^2+^. We did not investigate these GoSlo-SR-5-69 currents further that were produced by the *Xenopus* oocytes and were not associated with the expression of *Bma*-SLO-1F or *Ovo*-SLO-1A channels.

To determine the influence of extracellular Ca^2+^ on the GoSlo-SR-5-69 potentiation of emodepside, we compared results from oocytes expressing *Bma*-SLO-1F and *Ovo*- SLO-1A that were exposed to extracellular Ca^2+^ with oocytes that had Ca^2+^ replaced with equimolar Co^2+^. GoSlo-SR-5-69 still potentiated the emodepside responses in the absence of Ca^2+^ (Fig 7A, 7B, 7D and 7E). Although the mean currents appeared smaller in the absence of extracellular Ca^2+^, the differences in percentage increase in the emodepside currents by GoSlo-SR-5-69 were not significant (Fig 7C and F). These observations suggest that GoSlo-SR-5-69 potentiation is not mediated by entry of extracellular Ca^2+^ that could increase SLO-1 K channel opening.

**Fig 7.**
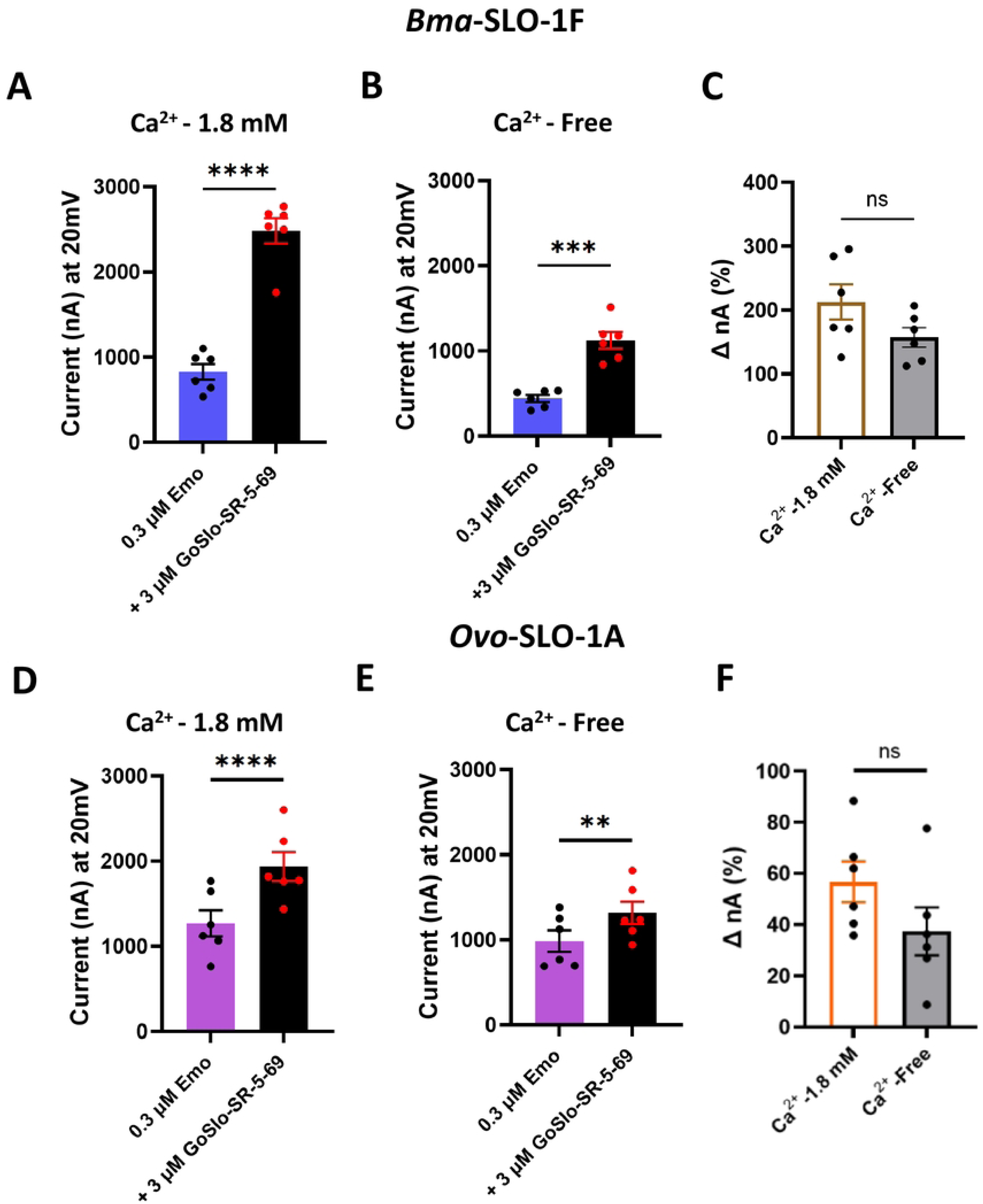
Effects of extracellular Ca^2+^ on GoSlo-SR-5-69 potentiation of *Bma*-SLO-1F and *Ovo*-SLO-1A-mediated emodepside responses. **A.** Bar chart (mean ± S.E.M) showing current (in nA) produced by *Bma*-SLO-1F expressing oocytes in response to 0.3 µM emodepside alone and in combination with 3 µM GoSlo-SR-5-69 in the presence of normal recording solution consisting of 1.8 mM added Ca^2+^. Blue bar: 0.3 µM emodepside alone (n = 6); Black bar: 3 µM GoSlo-SR-5-69 co-applied with 0.3 µM emodepside (n = 6). \*\*\*\**P* < 0.0001 significantly different as indicated; paired two-tailed student t-test **B.** Bar chart (mean ± S.E.M) showing current (in nA) produced by oocytes expressing the *Bma*-SLO-1F channel in response to 0.3 µM emodepside alone and in combination with 3 µM GoSlo-SR-5-69 in the absence of 1.8 mM added Ca^2+^. Blue bar: 0.3 µM emodepside alone (n = 6); Black bar: 3 µM GoSlo-SR-5-69 co-applied with 0.3 µM emodepside (n = 6). \*\*\**P* < 0.001, significantly different as indicated, paired two-tailed student t-test. **C.** Bar chart (mean ± S.E.M) of percentage (%) increase in currents produced by emodepside and GoSlo-SR-5-69 co-application for *Bma*-SLO-1F expressing oocytes perfused with normal recording solution, containing 1.8 mM added Ca^2+^ (White bar with brown border, n = 6), or modified recording solution lacking added Ca^2+^(grey bar with black border, n = 6). *P* > 0.05, no statistical significance (ns) as indicated, unpaired two-tailed student t- test. **D.** Bar chart (mean ± S.E.M) showing current (in nA) produced by oocytes expressing the *Ovo*-SLO-1A channel in response to 0.3 µM emodepside alone and in combination with 3 µM GoSlo-SR-5-69 in the presence of 1.8 mM added Ca^2+^. Pink bar: 0.3 µM emodepside alone (n = 6); Black bar: 3 µM GoSlo-SR-5-69 co-applied with 0.3 µM emodepside (n = 6). **E.** Bar chart (mean ± S.E.M) showing current (in nA) produced by *Ovo*-SLO-1A expressing oocytes in response to 0.3 µM emodepside alone and in combination with 3 µM GoSlo-SR-5-69 in the absence of 1.8 mM added Ca^2+^. Pink bar: 0.3 µM emodepside alone (n = 6); Black bar: 3 µM GoSlo-SR-5-69 co-applied with 0.3 µM emodepside (n = 6). \*\**P* < 0.0001 significantly different as indicated; paired two-tailed student t-test; \*\*\**P* < 0.01, significantly different as indicated, paired two-tailed student t- test. **F.** Bar chart (mean ± S.E.M) of percentage (%) increase in currents produced by emodepside and GoSlo-SR-5-69 co-application for *Ovo*-SLO-1A expressing oocytes in the presence of 1.8 mM added Ca^2+^ (White bar with orange border, n = 6), or modified recording solution lacking added Ca^2+^(Grey bar with black border, n = 6). *P* > 0.05, no statistical significance (ns) as indicated, unpaired two-tailed student t-test.

### GoSlo-SR-5-69 increases emodepside potency and efficacy for *Bma*-SLO-1F

Our observation of GoSlo-SR-5-69 potentiating emodepside responses in our previous experiments prompted us to investigate further the positive allosteric modulation. We compared the effect of GoSlo-SR-5-69 on the concentration-response relationships of emodepside on oocytes expressing the *Bma*-SLO-1F channel with the concentration effect of emodepside alone. For the experiments we alternated between oocytes tested with emodepside alone and those pre-treated with GoSlo-SR-5-69. A representative trace of emodepside current responses alone and in the presence of 3 µM GoSlo-SR-5- 69 is shown in Fig S2A&B.

Our analysis revealed a left shift in the sigmoidal concentration-response curve for emodepside in the presence of GoSlo-SR-5-69 (Fig 8A). The EC_50_ was 1.40 ± 0.15 µM and *R_max_* was 2679 ± 318 nA, n = 6, for emodepside in the absence of GoSlo-SR-5-69. The EC_50_ was 0.20 ± 0.02 µM and the R*_max_* value was 4493 ± 433 nA, n = 6 for emodepside in the presence of 3 µM GoSlo-SR-5-69, n = 6.

**Fig 8.**
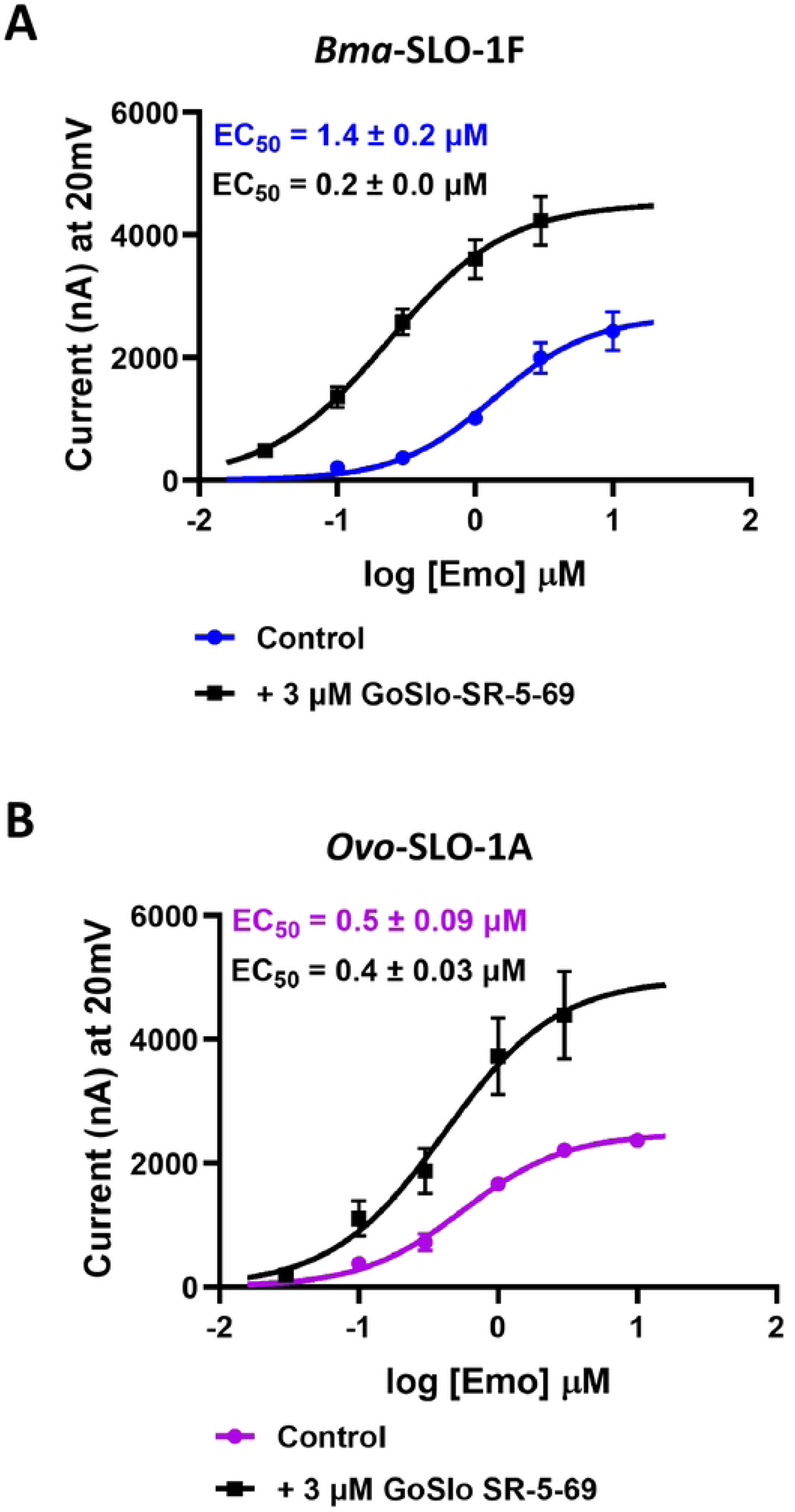
Effects of GoSlo-SR-5-69 as a positive allosteric modulator on *Bma*-SLO-1F and *Ovo*-SLO-1A channels on emodepside mediated response. **A.** Concentration-response plots for emodepside alone, (n = 6, blue) and emodepside in the presence of 3 µM GoSlo-SR-5-69 (n = 6, black) for the *Bma*-SLO-1F channel. **B.** Emodepside concentration-response plots for *Ovo*-SLO-1A in the presence of emodepside alone, (n = 6, pink) and emodepside in the presence of 3 µM GoSlo-SR-5-69 (n = 6, black). Bottom was constrained to zero for curve fitting.

The sensitivity of the *Bma*-SLO-1F channel was increased 7-fold to emodepside in the presence of 3 µM GoSlo-SR-5-69. Moreover, the 1.7-fold increase in *R_max_* also implies an increased efficacy of emodepside when GoSlo-SR-5-69 is present.

### GoSlo-SR-5-69 increases emodepside efficacy for *Ovo*-SLO-1A

The potentiation of emodepside responses were also seen for oocytes expressing the *Ovo*-SLO-1A channel. Using a similar approach as mentioned previously for *Bma*- SLO-1F, we investigated emodepside’s concentration-response relationships on *Ovo*- SLO-1A in the presence of 3 µM GoSlo-SR-5-69. A representative trace of emodepside current responses alone and in the presence of 3 µM GoSlo-SR-5-69 is shown in S3A and B Fig respectively.

The concentration response plots showed GoSlo-SR-5-69 is also a positive allosteric modulator of *Ovo*-SLO-1A (Fig 8B). The EC_50_ was 0.50 ± 0.09 µM and the *R_max_* was 2454 ± 87 nA, (n = 6), for emodepside in the absence of GoSlo-SR-5-69: the EC_50_ was 0.40 ± 0.03 µM and *R_max_* was 4928 ± 830 nA, (n = 6) in the presence of 3 µM GoSlo-SR-5-69. GoSlo-SR-5-69 did not cause a significant shift in EC_50_, but significantly increased the *R_max_*, confirming that GoSlo-SR-5-69 is also a positive allosteric modulator of the Ovo-SLO-1A receptor.

### Molecular docking proposes favorable binding mode of emodepside at S6 pocket in *Ovo*-SLO-1A and *Bma*-SLO-1F channels

Our molecular docking of emodepside in the SLO-1 K channel for both *B. malayi* and *O. volvulus* suggests that it adopts a similar pose, consistent with the findings for the cryo-EM structure of *D. melanogaster* SLO-1 K channel [15] (Fig 9A and B). The favorable hydrophobic pocket at the S6 site suitably accommodates emodepside, suggested by GLIDE to have very favorable docking scores, and an improvement on those of the *D. melanogaster* SLO-1 conformation, at −9.3 kcal/mol for *Ovo*-SLO-1A, and −9.7 kcal/mol for *Bma*-SLO-1F compared to −8.6 kcal/mol for *D. melanogaster*.

**Fig 9.**
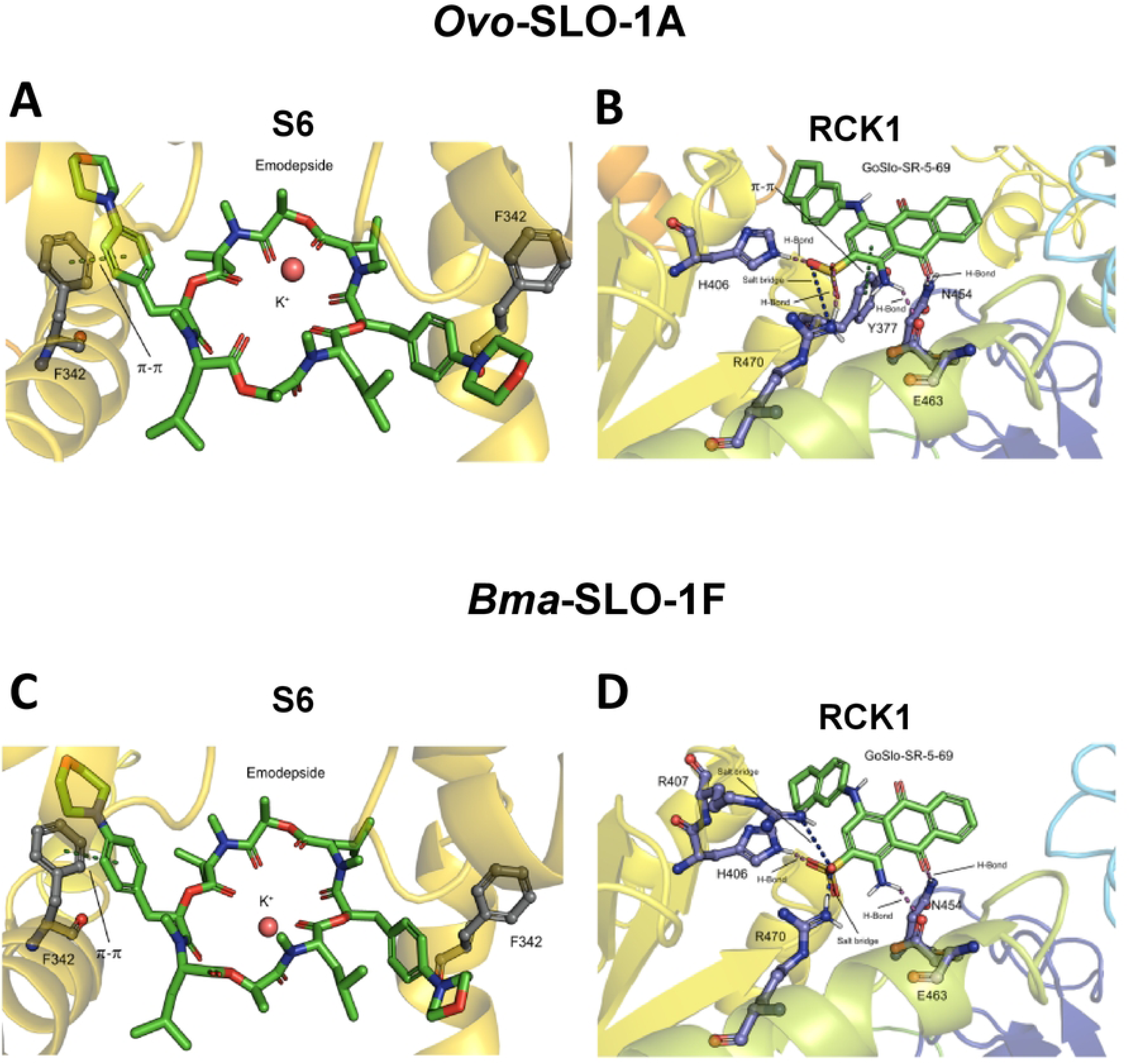
Identified poses of emodepside and GoSlo-SR-5-69 in *Ovo*-SLO-1A and *Bma*- SLO-1F channels. **A.** Emodepside bound at the S6 pocket for *Ovo*-SLO-1A channel. **B.** GoSlo-SR-5-69 bound in the RCK1 pocket for *Ovo*-SLO-1A channel. **C.** Emodepside bound at the S6 pocket for *Bma*-SLO-1F channel **D.** GoSlo-SR-5-69 bound at the RCK1 pocket for *Bma*-SLO-1F channel. For both channels, emodepside is bound below the selectivity filter, indicating the π-π stacking between F342 and emodepside. Displayed are also the stabilizing ligand-receptor interactions between GoSlo-SR-5-69 and the respective filarial nematode splice variant channels.

The top poses identified by GLIDE for both channels indicated a stabilizing π-π interaction between F342 and the phenyl rings of emodepside (Fig 9A and C), but these interactions were not found with the corresponding ligand F329 on the *D. melanogaster* binding pocket. After energy minimization, the RMSD difference in the emodepside structures between *Bma*-SLO-1F and *Ovo*-SLO-1A was low at 1.1 Å. The difference in the F342 residue position between the *Bma*-SLO-1F and *Ovo*-SLO-1A was 0.2 Å, with marginally better orientation between the phenyl rings for *Ovo*-SLO-1A. The RMSD between *D. melanogaster* and each of the nematode BK channels for the emodepside binding position was 6.5 Å for *Ovo*-SLO-1A and 6.4 Å for *Bma*-SLO-1F.

### Molecular docking suggests the RCK1 binding site for GoSlo-SR-5-69

To elucidate the potential positive allosteric modulation mechanism of GoSlo-SR- 5-69 in the SLO-1 K channels of *Ovo*-SLO-1A and *Bma*-SLO-1F, we probed the additional drug-binding pockets, RCK1 and RCK2 previously identified [15] using molecular docking. We identified binding poses for the GoSlo-SR-5-69 in both the RCK1 and RCK2 pockets; but the docking scores for the RCK1 pocket were much more favorable for both channels compared to those of the RCK2 pocket, with an average GlideScore difference of 2.4kcal/mol between two pockets. The GlideScores for the RCK1 pocket were −3.0 kcal/mol for *Bma*-SLO-1F and −2.5 kcal/mol for *Ovo*-SLO-1A. More non-covalent and π interactions are present within the RCK1 pocket, with the following key interactions between the ligand and channels for both *Bma*-SLO-1F and *Ovo*-SLO-1A: H-Bond (*Bma*- SLO-1F: H406, E463, N454, *Ovo*-SLO-1A: H406, R470, E463, N454), Salt bridge (*Bma*- SLO-1F: R407, R470, *Ovo*-SLO-1A: R470), π-π stacking (*Ovo*-SLO-1A: Y377) (Fig 9B and D). Overall, this suggests that the RCK1 pocket is the preferred binding site for GoSlo-SR-5-69 over RCK2.

## Discussion

### Emodepside is more potent on *Ovo*-SLO-1A than previously reported

Here we demonstrate that emodepside is more potent on *Ovo*-SLO-1A than *Bma*- SLO-1F. This differs from previous reports that suggest that the potency of emodepside was greater for *Bma*-SLO-1F than *Ovo*-SLO-1A [16]. The discrepancy in potency can be attributed to differences in our experimental techniques. First, emodepside is very lipophilic and difficult to wash off, as we observed in our preparations and reported previously [18, 19]. Hence, we conducted cumulative concentration-response experiments, eliminating wash steps between successive drug applications. Second, emodepside activation of SLO-1 K channels produces currents that are slow in onset, gradually increasing over a longer period of 5 minutes [19]. Consequently, we perfused emodepside for a longer duration to achieve plateau of the elicited currents during drug application. We point out that the expression levels of the *Ovo*-SLO-1A and *Bma*-SLO- 1F channels varied, affecting the amplitude of the current responses between individual oocytes and different batches of oocytes. To control this variation, we compared effects on the same batches of oocytes and calculated the mean current responses of six or more oocytes. Together, these alterations in protocol improved the estimation of emodepside potency on each channel isoform.

### Explanation of differences in *Bma*-SLO-1F and *Ovo*-SLO-1A sensitivity to emodepside

Our homology modelling revealed that emodepside can bind in the S6 pocket of the pore domain (PD) below the selectivity filter of *Ovo*-SLO-1A and *Bma*-SLO-1F, Fig 9A and C and Fig 10A and B. The PD is 100% identical between both channel isoforms and emodepside interacts with the same amino acid residues (S4 Fig). This suggests that the difference in emodepside potency should be attributed to allosteric effects of other structural domains such as the cytosolic tail domain (CTD) and voltage sensor domain (VSD) that modulate SLO-1 K channel function.

**Fig 10.**
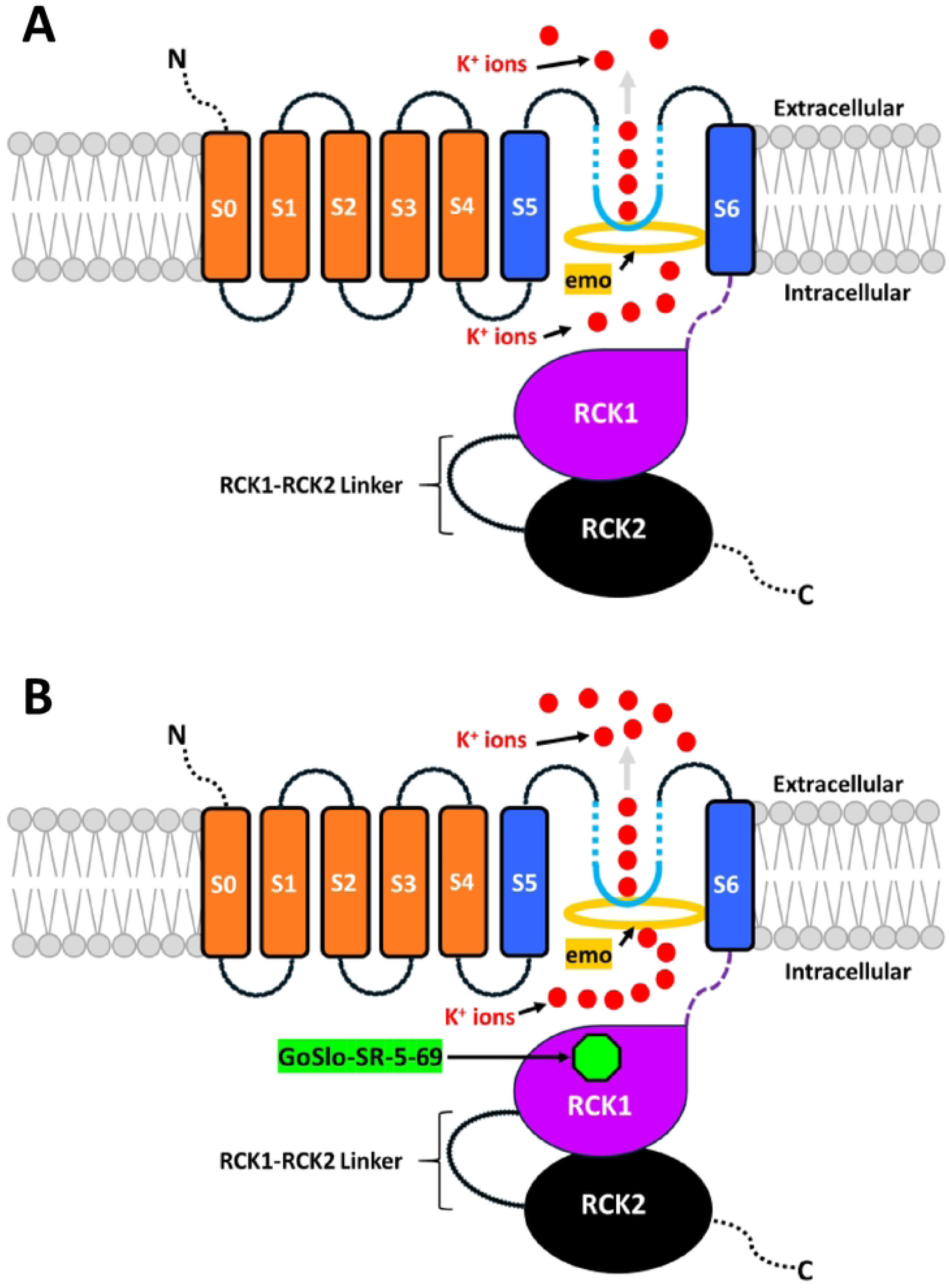
Summary diagram of the putative mechanism of GoSlo-SR-5-69 alone and in combination with emodepside on *Bma*-SLO-1F and *Ovo*-SLO-1A channels. **A.** Emodepside (gold ring) binds within the pore domain (PD) beneath the selectivity filter (light blue inverted block arc) and directly activates either *Bma*-SLO-1F or *Ovo*-SLO-1A channels resulting in the translocation of K^+^ ions (red circles) out of the cell (outward current). **B.** Activation of *Bma*-SLO-1F or *Ovo*-SLO-1A by emodepside (gold ring) in the PD. GoSlo-SR-5-69 (green octagon) then binds to a putative allosteric site (RCK1 domain) of the channel giving rise to stabilization of the channel gating ring in the open state. Subsequently, K^+^ permeation and translocation are further increased leading to greater current amplitude and the potentiation of emodepside response. The removal of GoSlo-SR-5-69 results in ion translocation almost back to normal levels previously seen in the presence of emodepside binding alone.

The CTD is connected to the C-terminus of the PD and serves as an intracellular sensor of Ca^2+^, Mg^2+^ and other intracellular ligands. In addition to its chemo-mechanical role, the CTD also possesses regulatory regions that impact SLO-1K channel function. We observed that oocytes expressing *Ovo*-SLO-1A channels produced higher currents and greater slope conductance’s than *Bma*-SLO-1F. Previous studies have reported a direct correlation between the membrane conductance and the number of ion channels expressed [20, 21]. This may be explained if: 1) there are more *Ovo*-SLO-1A channels expressed than *Bma*-SLO-1F channels; 2) the probability of the *Ovo*-SLO-1A channels being open may be higher than the *Bma*-SLO-1F channels or 3) the single channel conductance of the *Ovo*-SLO-1A channels are bigger than the *Bma*-SLO-1F channels.

An acidic cluster-like motif (DDXXDXXXI) trafficking signal within the intracellular RCK1-RCK2 linker of the CTD, has been implicated in the regulation of cell surface expression of human BK channels [22]. We identified this motif in the *Bma*-SLO-1F and *Ovo*-SLO-1A channel sequences, but with a glutamic acid E646 residue instead of aspartic acid (D), (EDXXDXXXI) as shown in red font and highlighted in turquoise in Fig S4. Additionally, two amino acid residues of the motif sequence for *Ovo*-SLO-1A (EDIIDVSLI) and *Bma*-SLO-1F (EDIMDISLI) were not the same. Based on the acidic cluster-like motif sequence differences here that affect cell surface expression, we suggest that *Ovo*-SLO-1A channels are exported in higher numbers to the oocyte membrane, than *Bma*-SLO-1F. Hence, the smaller effect of emodepside on *Bma*-SLO- 1F channels may be attributed to reduced ion channel expression. If a similar limiting cell surface regulation mechanism is adopted in the adult *B. malayi* parasite, it could be part of the reason *B. malayi* is less sensitive to emodepside making it the dose-limiting filarial nematode species.

In addition to SLO-1 K channel expression, variation in the tension of the RCK1- RCK2 loop in the CTD influences SLO-1 K activation. We found that the RCK1-RCK2 loop for *Bma*-SLO-1F was shorter than that of *Ovo*-SLO-1A with a total length of 87 and 102 amino acids, respectively. Mammalian SLO-1 K channels generally have at least 101 amino acids separating the two RCK domains. Reducing the length of the RCK1-RCK2 loop affects SLO-1 K channel function and consequently macroscopic currents [23]. A minimum length of approximately 70 amino acids is required in the connecting loop for the channel to be functional [23]. This shorter loop for *Bma*-SLO-1F is predicted to reduce the channel openings due to a stronger tension on the gate that is needed for openings. The shorter amino acid length separating the RCK1-RCK-2 loop is predicted to reduce the overall lower currents for *Bma*-SLO-1F produced in the absence and presence of emodepside.

Differences in emodepside sensitivity between *Bma*-SLO-1F and *Ovo*-SLO-1A may also be attributed to non-conserved amino acid residues in the VSD. The PD is surrounded by four VSDs that forge strong electromechanical coupling [24, 25]. Changes in membrane potential induces conformational changes in the central cavity of the channel, resulting in its activation or deactivation [26, 27]. We identified 11 amino acid residue differences in the VSD of *Ovo*-SLO-1A and *Bma*-SLO-1F. Despite these differences in the VSD, our voltage step analysis showed that emodepside produced increased currents for both channel isoforms in the absence of increased intracellular Ca^2+^ and hyperpolarized potentials. This provides further confirmation that emodepside deregulates voltage and Ca^2+^ sensitivity of SLO-1 K channels [15]. Nevertheless, the elicited emodepside currents were inhibited at potentials exceeding +20 mV for *Bma*- SLO-1F, in a manner like that seen for the *Caenorhabditis elegans* SLO-1A channel [18]. The non-conserved residues in the VSD may alter the structural architecture of the VSD, thus influencing interactions with the PD during channel pore activation. However, further studies are needed to confirm these assumptions.

### The significance and mode of action of the potentiator, GoSlo-SR-5-69

Drug combination therapies have been widely used for the treatment of malaria, tuberculosis, and HIV [28–33]. Despite the therapeutic potential of emodepside for the treatment of human onchocerciasis, its use as an adulticidal treatment of lymphatic filariasis has not been pursued due to reduced emodepside potency on *Brugia spp.* [6, 34]. Increasing the potency of emodepside on the less sensitive *Brugia spp*. could offer a significant therapeutic advance.

Putative candidates for drug combination and toxicity testing are the negatively charged activators (NCAs) like GoSlo-SR-5-69. The GoSlo-SR family of compounds have been reported to activate mammalian SLO-1 K channels through interactions with amino acid residues in the transmembrane domain [14, 35]. Additionally, their negatively charged sulphonate group has been predicted to attract numerous K^+^ ions to the channel pore, increasing ion occupancy and consequently channel conductance, based on the mechanism of other NCAs [36]. We found however in our experiments that GoSlo-SR-5- 69 does not activate the channels of *Bma*-SLO-1F or *Ovo*-SLO-1A. Nevertheless, we observed very clear effects of GoSlo-SR-5-69 in combination with emodepside, that produced emodepside potentiation. Remarkably, the sensitivity of *Bma*-SLO-1F to emodepside, measured by its EC_50_ was increased 7-fold, while the EC_50_ of *Ovo*-SLO-1A remained unchanged, although there was an increase in the *R_max_*.

Our homology modelling revealed that GoSlo-SR-5-69 can bind to the RCK1 pocket of both channels, at the Rossmann-fold subdomain (βA-βF) in the Ca^2+^ bound open state. The Rossmann-fold subdomain is known to form the central core of the gating ring [25]. Therefore, we propose a mechanism for emodepside potentiation that involves the binding of emodepside at the pore domain, leading to channel activation. Access of GoSlo-SR-5-69 to the channel in its open conformation allows favorable binding of the molecule at the RCK1 domain where it stabilizes the open-state of the channel, thereby potentiating emodepside response. GoSlo-SR-5-69 would not have access to this binding site in the closed conformation and by itself does not open the channel.

We also note that the greater level of emodepside potentiation by GoSlo-SR-5-69 on *Bma*-SLO-1F could be attributed to minor differences in GoSlo-SR-5-69 interactions with the channels. GoSlo-SR-5-69 was found to dock to the same residues except for R407 and P409 for *Ovo*-SLO-1A (S4 Fig). Hence, slight differences in binding sites may influence the degree of potentiation.

In conclusion, our data has highlighted the pharmacological diversity of two highly conserved filarial nematode SLO-1 K channels. We demonstrate that both *Bma*-SLO-1F and *Ovo*-SLO-1A are activated by emodepside. *Bma*-SLO-1F is less sensitive to emodepside than *Ovo*-SLO-1A, providing an explanation for the lack of emodepside efficacy on *Brugia spp.* in contrast to *Onchocerca spp*. We also show that GoSlo-SR-5- 69 is not an activator of the splice variant channels but acts as a positive allosteric modulator: 1) increasing the potency and efficacy of emodepside on *Bma*-SLO-1F channels; and 2) increasing emodepside efficacy on *Ovo*-SLO-1A. The identification of emodepside binding site and its unique π-π interaction with F342 highlights a putative nematode-specific interaction that facilitates emodepside selectivity. Additionally, predicting the GoSlo-SR-5-69 most favorable binding site on the splice variant channels provides a proposed mechanism of action for GoSlo-SR-5-69 potentiation of emodepside. We provide a mechanism for a strategy to increase emodepside potency on a SLO-1 K splice variant isoform for *B. malayi* through drug combination.

## Methods

### Sequence Analysis

*Bma*-SLO-1F and *Ovo*-SLO-1A amino acid sequences were acquired from the *B. malayi* and *O. volvulus* genome using WormBase ParaSite (parasite.wormbase.org). Sequence alignment was conducted using EMBOSS Needle pairwise sequence alignment tools, with EBLOSUM62 matrix, a default gap penalty of 10 and extension penalty of 0.5 [37], to determine sequence identity and similarity between both species and isoforms.

Sequence annotation was achieved using previously published alignment information by highlighting the voltage sensor domain (VSD), pore domain (PD) and the regulator of potassium conductance domains (RCK1 and RCK2) [15]. To further estimate conservation of each individual domain between isoforms, alignments and sequence identity analysis were also conducted for each domain individually.

### Cloning of Brugia malayi slo-1f and Onchocerca volvulus slo-1a

Primers for *Brugia malayi slo-1f* and *Onchocerca volvulus slo-1a* isoforms were designed with sequences flanking the pT7TS-rich expression vector that included the restriction site (NheI). PCR amplification was conducted on *B. malayi slo-1f* that was previously cloned in the pCDNA3.1 vector. In contrast, *Onchocerca volvulus slo-1a* was synthesized by Life Technologies GeneArt. Subsequently, both amplicons were separated on a 1% Agarose SYBR Safe gel, purified using NucleoSpin Gel and PCR Clean-up Kit (Macherey-Nagel) and cloned into the pT7TS-rich vector by using Infusion HD Cloning Kit (Takara Bio USA, Inc) according to the manufacturer’s protocols. Once cloned, the plasmids were verified by sequencing.

### In vitro transcription of Bma-slo-1f and Ovo-slo-1a

The pT7TS-rich plasmids containing cloned products of either *Bma-slo-1f* or *Ovo-Slo-1a* were linearized by SmaI and BamHI respectively and purified. Capped cRNAs were then synthesized from the linearized vectors containing the *B. malayi* and *O. volvulus Slo-1* isoforms previously mentioned using the T7 mMessage mMachine Kit (Ambion, USA). The cRNAs were stored at −80°C until further use.

### Heterologous expression of *Bma*-SLO-1F and *Ovo*-SLO-1F receptors in *Xenopus laevis* oocytes

Defolliculated *Xenopus laevis* oocytes were purchased from Ecocyte Bioscience (Austin, TX, USA) and Xenopus 1 Corp (Dexter, MI, USA). Heterologous expression of the *Bma*-SLO-1F receptor was achieved by injecting 15 ng of cRNA in a total volume of 50 nL in nuclease-free water. Each oocyte was microinjected into the cytoplasm of the animal pole region using a Drummond Nanoject II microinjector (Drummond Scientific, Broomall, PA, USA). After injection, oocytes were incubated at 17°C in a sterile 96-well culture plate containing 300 μl of incubation solution (100 mM NaCl, 2 mM KCl, 1.8 mM CaCl_2_.2H_2_O, 1 mM MgCl_2_.6H_2_O, 5 mM HEPES, 2.5 mM Na pyruvate, 100 U/mL penicillin and 100 μg/mL streptomycin, pH 7.5) in each well. Incubation solution was changed daily during the period of incubation. The same procedure was also conducted for the *Ovo*- SLO-1A receptor. Experiments were performed on oocytes within 5 – 6 days post injection.

### Two-microelectrode voltage clamp (TEVC) electrophysiology

TEVC was conducted at room temperature by impaling oocytes with two microelectrodes; a current injecting electrode, Im, used to inject the required current for holding the membrane at a set voltage, and a voltage sensing electrode, Vm. The microelectrodes were pulled using a Flaming/Brown horizontal electrode puller (Model P- 97; Sutter Instruments, Novato, CA, USA) and filled with 3 M KCl. Each electrode tip was broken with a piece of Kimwipe paper (Kimtech Science^TM^, Fisher) to achieve a resistance of 2 – 5 MΏ in recording solution (88 mM NaCl, 2.5 mM KCl, 1 mM MgCl_2_.6H_2_O, 1.8 mM CaCl_2_.2H_2_O and 5 mM HEPES, at pH 7.4). To investigate the concentration-response relationship of emodepside on the expressed *Bma*-SLO-1F or *Ovo*-SLO-1A receptors, oocytes were voltage clamped at a steady-state potential of +20 mV with an Axoclamp 2B amplifier (Molecular Devices, Sunnyvale, CA, USA). Amplified signals were converted from analog to digital format by a Digidata 1322A digitizer (Molecular Devices, CA, USA) and all data were acquired on a desktop computer with the Clampex 10.3 data acquisition software (Molecular Devices, Sunnyvale, CA, USA). In addition, the same protocol was also used to test the effects of GoSlo-SR-5-69 alone or in combination with emodepside.

### Voltage Step Electrophysiology

Voltage step experiments were conducted using the two-electrode voltage-clamp technique to determine current-voltage relationships of each receptor in the absence and presence of 1 µM emodepside. Briefly, oocytes expressing either *Bma*-SLO-1F or *Ovo*- SLO-1A channels were impaled and subjected to a current-voltage protocol that consisted of 500 ms voltage steps from −120 to + 60 mV in 20 mV increments, starting from a holding position of −70 mV for 1 s between each step. Plateau currents were recorded at a frequency of 5000 Hz during clamping and perfused with recording solution: (88 mM NaCl, 2.5 mM, KCl, 1 mM MgCl_2_.6H_2_O, 1.8 mM CaCl_2_.2H_2_O and 5 mM HEPES, at pH 7.4). The results of the voltage steps were evaluated and analyzed using the ClampFit 10.3 software (Molecular Devices), whereas current-voltage curves (IVCs) were prepared using Graphpad Prism 10.1.1(GraphPad Software, Inc., USA).

### Chemicals

Emodepside was purchased from Advanced ChemBlock Inc (Hayward, CA, USA). GoSlo-SR-5-69 was purchased from Tocris Bioscience (Bristol, UK). Stock solutions of emodepside were prepared at 0.1, 0.3, 1, 3, 10 and 30 mM in dimethyl sulfoxide (DMSO) solutions prior to experimentation, then diluted in recording solution. Stock solutions of GoSlo-SR-5-69 were made in DMSO, at 50 mM, then diluted in recording solution to the required concentrations (3 µM). The final DMSO concentration did not exceed 0.1% in the experimental solutions.

### Drug application

Emodepside is known to be lipophilic, thus making it difficult to wash off completely from the *Xenopus laevis* oocyte preparation after application. Additionally, emodepside concentrations exceeding 10 µM showed evidence of precipitation and a limit of solubility. To estimate EC_50_ values, we utilized a cumulative concentration-response protocol (no wash steps between drug application) and a maximum concentration of 10 µM emodepside. For our recordings, the times for drug applications were selected to allow the currents recorded to reach a stable plateau. At the beginning of experiments the *Xenopus* oocytes were perfused with drug free recording solution for 1 min, to obtain stable initial resting currents. For our drug applications we used successive applications of increasing concentrations of emodepside (0.1 – 10 µM) for 5 mins per concentration. Washing of the *Xenopus* oocytes then followed for 3 mins.

The effects of the mammalian BK channel blocker, verruculogen was tested on *Bma*-SLO-1F and *Ovo*-SLO-1A expressing oocytes in a manner before and after the application of emodepside. Briefly, oocytes were perfused for 1 min with recording solution which was then followed by 2 mins application of verruculogen and subsequently application of emodepside for 5 mins. These experiments were repeated in the opposite order where emodepside was applied first for 5 mins, followed by verruculogen for 2 mins.

To investigate the effects of GoSlo-SR-5-69 (a mammalian BK channel activator), on *Bma*-SLO-1F and *Ovo*-SLO-1A channels, oocytes were perfused with recording solution for 1 min followed by application of 3 µM GoSlo-SR-5-69 for 5 mins and subsequently 0.3 µM emodepside (positive control) for 5 mins with a final wash step of recording solution for 3 mins.

To determine the effects of GoSlo-SR-5-69 in combination with emodepside, we employed a protocol for oocytes that involved 1 min perfusion of recording solution to obtain the control current levels. This was followed by the application of 0.3 µM emodepside for 5 mins, then co-application of 3 µM GoSlo-SR-5-69 in the continued presence of 0.3 µM emodepside for 5 mins and immediate wash off with 0.3 µM emodepside for 5 mins and a final wash with recording solution for 3 mins.

Finally, to evaluate the effects of 3µM GoSlo-SR-5-69 on emodepside concentration-response relationships, recording solution was applied for 1 min to each oocyte, followed by 3 µM GoSlo-SR-5-69 until evoked currents were stabilized. Next, 5 mins applications of increasing concentrations of emodepside (0.1 – 10 µM) were perfused in the continued presence of 3 µM GoSlo-SR-5-69. A 3 min wash off time was allowed at the end of the final concentration of emodepside.

### Homology Modelling and Molecular Docking

SLO-1 channels for *O. volvulus* and *B. malayi* were constructed from FASTA sequences obtained from the WormBase genome projects (*Bma*-SLO-1F and *Ovo*-SLO- 1A). Homology models were constructed using SWISS-MODEL, with the *Drosophila melanogaster* SLO channels in Ca^2+^ bound state (RCSB: 7PXE) as template [15]. The structure of emodepside was obtained from PubChem (CID6918632), and pre-docking structures of the channels were energy-minimized using GROMACS 2023.2, solvated in water and neutralized with potassium (K^+^) ions [38]. AMBER99SB-ILDN force field was used, and water molecules were parametrized with SPC/E [39]. Emodepside was parameterized with GAFF using Antechamber version 17.3 [40, 41]. The structure of GoSlo-SR-5-69 was obtained from PubChem (CID56944133), and prepared for docking using LigPrep in the Schrödinger package version 2023-4 at pH 7.0, with OPLS4 FF.

Docking was performed using GLIDE module in Schrödinger, docked to the homology models of SLO-1A for *O. volvulus*, SLO-1F for *B. malayi* and the cryo-EM structure for *D. melanogaster* SLO [42]. Docking for all structures was performed at Extended Precision (XP) level. The S6 emodepside pocket was defined as the position emodepside adopted in the energy-minimized structure. The RCK1 binding site was defined as the centroid of G430, M433, Y377, D411 for *O. volvulus* and *B. malayi* and for the RCK2, L777, H491, Y490, R781 for *B. malayi* and L792, H491, Y490, and R796 for *O. volvulus*.

Analysis of docking results was performed in the Schrödinger Maestro suite to identify non-covalent interactions between ligand and receptor, RMSD was calculated by aligning the structures in PyMOL (v.2.5.4).

### Data analysis

Our emodepside concentration-response experiments involved the use of Clampfit 10.3 (Molecular Devices, Sunnyvale, CA, USA) to measure peak current responses for each drug concentration (0.1 - 10 µM) per oocyte. GraphPad Prism 10.1.1 software (GraphPad Software Inc., USA) was used to generate Concentration-response curves using the log agonist vs. response equation (variable slope) to estimate EC_50_, *R_max_* and Hillslope (n*H*) values for both *Bma*-SLO-1F and *Ovo*-SLO-1A channels. We also used the unpaired two-tailed Student’s t-test to test for statistical significance. A p value < 0.05 was deemed significant. The analyzed results were expressed as mean ± S.E.M.

To obtain current-voltage curves (IVCs) from our voltage steps experiments, plateau currents elicited at each step potential (−120 to + 60 mV) were acquired for individual oocytes in Clampfit 10.3 (Molecular Devices, Sunnyvale, CA, USA). Mean currents for all replicate oocytes were plotted against their corresponding voltage step potentials to obtain IVCs using Graphpad Prism 10.1.1 software (GraphPad Software, Inc., USA). Mean currents between each treatment group of oocytes were compared for each step potential using two-way ANOVA and Tukey’s multiple comparison test to test for significance. To obtain and compare conductance changes in the absence and presence of 1 µM emodepside for *Bma*-SLO-1F and *Ovo*-SLO-1A channels, IVCs were analyzed for slopes between step potentials of −120 and +60 mV using linear regression analysis in Graphpad Prism 10.1.1(GraphPad Software, Inc., USA). Subsequently, statistical differences for slope values among treatment groups were analyzed using ANOVA, followed by the Tukey multiple comparison post-hoc test. Results were expressed as mean ± S.E.M.

To determine statistical significance of emodepside potentiation by GoSlo-SR-5- 69, we measured the mean currents evoked by 0.3 µM emodepside alone and compared it to the subsequent application of 3 µM GoSlo-SR-5-69 in combination with 0.3 µM emodepside on each oocyte using the unpaired two-tailed Student’s t-test in GraphPad Prism 10.1.1. Similar analyses were also conducted for the effect of extracellular Ca^2+^ on GoSlo-SR-5-69 potentiation of emodepside response involving recordings conducted in normal recording solution (1.8 mM added Ca^2+^) and modified recording solution (Ca^2+^-free). The results were expressed as the mean ± S.E.M.

Analysis involving the determination of GoSlo-SR-5-69 effects on *Bma*-SLO-1F and *Ovo*-SLO-1A mediated emodepside concentration-response relationship were conducted in a similar manner as previously described for the emodepside concentration-response experiments. Student’s *t-tests* were also used for comparing EC_50s_ and *R_max_* for emodepside alone and emodepside in combination with GoSlo-SR-5-69 using GraphPad Prism 10.1.1 software (GraphPad Software, Inc., USA). Results were expressed as the mean ± S.E.M.

## Acknowledgements

We express sincere gratitude to the Iowa State University DNA Facility for sequencing our cloned genes. Special thanks also to Dr. Michael Cho for the use of his nanodrop spectrophotometer machine.

## Author Contributions

**Conceptualization:** Mark McHugh, Sudhanva Kashyap, Alan P. Robertson, Richard J. Martin

**Formal Analysis:** Mark McHugh and Charity N Njeshi.

**Funding Acquisition**: Richard J Martin

**Investigation:** Mark McHugh, Charity N Njeshi, Nathaniel Smith, Han Sun

**Methodology:** Mark McHugh, Sudhanva Kashyap, Real Datta, Alan P. Robertson, Nathaniel Smith, Han Sun, Richard J. Martin.

**Project Administration:** Mark McHugh, Sudhanva Kashyap, Alan P. Robertson Nathaniel Smith, Han Sun, Richard J. Martin.

**Resources:** Han Sun, Richard J. Martin.

**Supervision:** Mark McHugh, Sudhanva Kashyap, Alan P. Robertson, Han Sun, Richard J. Martin.

**Validation:** Mark McHugh, Richard J. Martin

**Visualization:** Mark McHugh

**Writing – original draft:** Mark McHugh, Nathaniel Smith, Han Sun, Richard J. Martin.

**Writing – review & editing:** Mark McHugh, Nathaniel Smith, Han Sun, Richard J. Martin.

## Conflict Of Interest Statement

There are no conflicts of interest.

## Data Availability

The data from this study is available upon request from the corresponding author.

## Declaration of Transparency and Scientific Rigor

This Declaration acknowledges that this paper adheres to the principles for transparent reporting and scientific rigor of preclinical research as stated in the BJP Guidelines for Design and Analysis as recommended by funding agencies, publishers and other organizations engaged with supporting research.

## Supporting information

**Fig S1. Effects of GoSlo-SR-5-69 on *Bma*-SLO-1F and *Ovo*-SLO-1A channels in the absence of extracellular Ca^2+^. A.** Representative trace of water injected oocytes (n = 6). **B.** Representative trace of oocytes expressing the *Bma*-SLO-1F channel (n = 6). **C.** Representative trace of *Ovo*-SLO-1A expressing channel (n = 6).

**Fig S2. Effects of GoSlo-SR-5-69 on *Bma*-SLO-1F-mediated emodepside responses. A.** Representative current traces for two-electrode voltage-clamp recording showing outward currents for *Bma*-SLO-1F in response to increasing concentrations of emodepside (0.1 to 10 µM) at a steady-state holding potential of +20 mV. **B.** Representative current traces for two-electrode voltage-clamp recording showing outward currents for *Bma*-SLO-1F in response to increasing concentrations of emodepside (0.1 to 10 µM) in the presence of 3 µM GoSlo-SR-5-69 at a holding potential of +20 mV.

**Fig S3. Effects of GoSlo-SR-5-69 *Ovo*-SLO-1A-mediated emodepside responses. A.** Representative current traces for two-electrode voltage-clamp recording showing outward currents for *Ovo*-SLO-1A in response to increasing concentrations of emodepside (0.1 to 10 µM) at a steady-state holding potential of +20 mV. **B.** Representative current traces for two-electrode voltage-clamp recording showing outward currents for *Ovo*-SLO-1A in response to increasing concentrations of emodepside (0.1 to 10 µM) in the presence of 3 µM GoSlo-SR-5-69 at a holding potential of +20 mV.

**Fig S4. Amino acid sequence alignment of *Bma*-SLO-1F and *Ovo*-SLO-1A.** The voltage sensor domain (VSD; orange boxes), pore gate domain (PD; blue boxes) comprising the selectivity filter (light blue box), and two C-terminal domains for regulator of K^+^ conductance (RCK1; pink boxes and RCK2; black boxes) are indicated. Amino acids which are not identical between filarial species are highlighted by a black background. Gaps are indicated by “_” symbols for amino acid residues that are missing. Residues that are predicted to be involved in emodepside binding are highlighted by a yellow background in the PD. Putative amino acid residues involved in GoSlo-SR-5-69 binding are highlighted by a light green background in the RCK1 domain. Note that both *Bma*-SLO-1F and *Ovo*-SLO-1A have conserved amino acids interacting with GoSlo-SR-5-69 except for R407 and P409 that are not involved in binding for *Ovo*-SLO-1A. The acidic cluster-like motif (EDXXDXXXI) trafficking signal residues (underlined aqua) for *Bma*-SLO-1F and *Ovo*-SLO-1A is also shown in red font.

## Notes

### Competing Interest Statement

The authors have declared no competing interest.

## References

1. Taylor MJ, Hoerauf A, Bockarie M. Lymphatic filariasis and onchocerciasis. Lancet. 2010;376(9747):1175–85. Epub 20100823. doi: 10.1016/S0140-6736(10)60586-7. PubMed PMID: 20739055.

2. WHO. Global programme to eliminate lymphatic filariasis: progress report, 2022. 2023; 41(98), 489–502.

3. Omura S, Crump A. Ivermectin: panacea for resource-poor communities? Trends Parasitol. 2014;30(9):445–55. Epub 20140812. doi: 10.1016/j.pt.2014.07.005. PubMed PMID: 25130507.

4. Critchley J, Addiss D, Ejere H, Gamble C, Garner P, Gelband H, et al. Albendazole for the control and elimination of lymphatic filariasis: systematic review. Trop Med Int Health. 2005;10(9):818–25. doi: 10.1111/j.1365-3156.2005.01458.x. PubMed PMID: 16135187.

5. Hawking F. A review of progress in the chemotherapy and control of filariasis since 1955. Bull World Health Organ. 1962;27(4-5):555–68. PubMed PMID: 13953210; PubMed Central PMCID: PMCPMC2555874.

6. Hübner MP, Townson S, Gokool S, Tagboto S, Maclean MJ, Verocai GG, et al. Evaluation of the in vitro susceptibility of various filarial nematodes to emodepside. Int J Parasitol Drugs Drug Resist. 2021;17:27–35. Epub 20210728. doi: 10.1016/j.ijpddr.2021.07.005. PubMed PMID: 34339934; PubMed Central PMCID: PMCPMC8347670.

7. Kashyap SS, Verma S, Voronin D, Lustigman S, Kulke D, Robertson AP, et al. Emodepside has sex-dependent immobilizing effects on adult Brugia malayi due to a differentially spliced binding pocket in the RCK1 region of the SLO-1 K channel. PLoS Pathog. 2019;15(9):e1008041. Epub 20190925. doi: 10.1371/journal.ppat.1008041. PubMed PMID: 31553770; PubMed Central PMCID: PMCPMC6779273.

8. Bull K, Cook A, Hopper NA, Harder A, Holden-Dye L, Walker RJ. Effects of the novel anthelmintic emodepside on the locomotion, egg-laying behaviour and development of Caenorhabditis elegans. Int J Parasitol. 2007;37(6):627–36. Epub 20061127. doi: 10.1016/j.ijpara.2006.10.013. PubMed PMID: 17157854.

9. Guest M, Bull K, Walker RJ, Amliwala K, O’Connor V, Harder A, et al. The calcium-activated potassium channel, SLO-1, is required for the action of the novel cyclo-octadepsipeptide anthelmintic, emodepside, in Caenorhabditis elegans. Int J Parasitol. 2007;37(14):1577–88. Epub 20070521. doi: 10.1016/j.ijpara.2007.05.006. PubMed PMID: 17583712.

10. Welz C, Krüger N, Schniederjans M, Miltsch SM, Krücken J, Guest M, et al. SLO-1- channels of parasitic nematodes reconstitute locomotor behaviour and emodepside sensitivity in Caenorhabditis elegans slo-1 loss of function mutants. PLoS Pathog. 2011;7(4):e1001330. Epub 20110407. doi: 10.1371/journal.ppat.1001330. PubMed PMID: 21490955; PubMed Central PMCID: PMCPMC3072372.

11. Assmus F, Hoglund RM, Monnot F, Specht S, Scandale I, Tarning J. Drug development for the treatment of onchocerciasis: Population pharmacokinetic and adverse events modeling of emodepside. PLoS Negl Trop Dis. 2022;16(3):e0010219. Epub 20220310. doi: 10.1371/journal.pntd.0010219. PubMed PMID: 35271567; PubMed Central PMCID: PMCPMC8912909.

12. Gillon JY, Dennison J, van den Berg F, Delhomme S, Dequatre Cheeseman K, Peña Rossi C, et al. Safety, tolerability and pharmacokinetics of emodepside, a potential novel treatment for onchocerciasis (river blindness), in healthy male subjects. Br J Clin Pharmacol. 2021;87(10):3949–60. Epub 20210331. doi: 10.1111/bcp.14816. PubMed PMID: 33759250; PubMed Central PMCID: PMCPMC8518114.

13. Krücken J, Holden-Dye L, Keiser J, Prichard RK, Townson S, Makepeace BL, et al. Development of emodepside as a possible adulticidal treatment for human onchocerciasis-The fruit of a successful industrial-academic collaboration. PLoS Pathog. 2021;17(7):e1009682. Epub 20210722. doi: 10.1371/journal.ppat.1009682. PubMed PMID: 34293063; PubMed Central PMCID: PMCPMC8297762.

14. Roy S, Large RJ, Akande AM, Kshatri A, Webb TI, Domene C, et al. Development of GoSlo-SR-5-69, a potent activator of large conductance Ca2+-activated K+ (BK) channels. Eur J Med Chem. 2014;75:426–37. Epub 20140203. doi: 10.1016/j.ejmech.2014.01.035. PubMed PMID: 24561672.

15. Raisch T, Brockmann A, Ebbinghaus-Kintscher U, Freigang J, Gutbrod O, Kubicek J, et al. Small molecule modulation of the Drosophila Slo channel elucidated by cryo-EM. Nat Commun. 2021;12(1):7164. Epub 20211209. doi: 10.1038/s41467-021-27435-w. PubMed PMID: 34887422; PubMed Central PMCID: PMCPMC8660915.

16. Bah GS, Schneckener S, Hahnel SR, Bayang NH, Fieseler H, Schmuck GM, et al. Emodepside targets SLO-1 channels of Onchocerca ochengi and induces broad anthelmintic effects in a bovine model of onchocerciasis. PLoS Pathog. 2021;17(6):e1009601. Epub 20210602. doi: 10.1371/journal.ppat.1009601. PubMed PMID: 34077488; PubMed Central PMCID: PMCPMC8202924.

17. Horrigan FT, Aldrich RW. Coupling between voltage sensor activation, Ca2+ binding and channel opening in large conductance (BK) potassium channels. J Gen Physiol. 2002;120(3):267-305. doi: 10.1085/jgp.20028605. PubMed PMID: 12198087; PubMed Central PMCID: PMCPMC2229516.

18. Kulke D, von Samson-Himmelstjerna G, Miltsch SM, Wolstenholme AJ, Jex AR, Gasser RB, et al. Characterization of the Ca2+-gated and voltage-dependent K+- channel Slo-1 of nematodes and its interaction with emodepside. PLoS Negl Trop Dis. 2014;8(12):e3401. Epub 20141218. doi: 10.1371/journal.pntd.0003401. PubMed PMID: 25521608; PubMed Central PMCID: PMCPMC4270693.

19. Buxton SK, Neveu C, Charvet CL, Robertson AP, Martin RJ. On the mode of action of emodepside: slow effects on membrane potential and voltage-activated currents in Ascaris suum. Br J Pharmacol. 2011;164(2b):453–70. doi: 10.1111/j.1476-5381.2011.01428.x. PubMed PMID: 21486286; PubMed Central PMCID: PMCPMC3188918.

20. Schulz DJ, Goaillard JM, Marder E. Variable channel expression in identified single and electrically coupled neurons in different animals. Nat Neurosci. 2006;9(3):356–62. Epub 20060129. doi: 10.1038/nn1639. PubMed PMID: 16444270.

21. Veys K, Labro AJ, De Schutter E, Snyders DJ. Quantitative single-cell ion-channel gene expression profiling through an improved qRT-PCR technique combined with whole cell patch clamp. J Neurosci Methods. 2012;209(1):227–34. Epub 20120620. doi: 10.1016/j.jneumeth.2012.06.008. PubMed PMID: 22728251.

22. Chen L, Jeffries O, Rowe IC, Liang Z, Knaus HG, Ruth P, et al. Membrane trafficking of large conductance calcium-activated potassium channels is regulated by alternative splicing of a transplantable, acidic trafficking motif in the RCK1-RCK2 linker. J Biol Chem. 2010;285(30):23265–75. Epub 20100517. doi: 10.1074/jbc.M110.139758. PubMed PMID: 20479001; PubMed Central PMCID: PMCPMC2906319.

23. Lee JH, Kim HJ, Kim HD, Lee BC, Chun JS, Park CS. Modulation of the conductance-voltage relationship of the BK(Ca) channel by shortening the cytosolic loop connecting two RCK domains. Biophys J. 2009;97(3):730–7. doi: 10.1016/j.bpj.2009.04.058. PubMed PMID: 19651031; PubMed Central PMCID: PMCPMC2718151.

24. Sun L, Horrigan FT. A gating lever and molecular logic gate that couple voltage and calcium sensor activation to opening in BK potassium channels. Sci Adv. 2022;8(50):eabq5772. Epub 20221214. doi: 10.1126/sciadv.abq5772. PubMed PMID: 36516264; PubMed Central PMCID: PMCPMC9750137.

25. Yang H, Zhang G, Cui J. BK channels: multiple sensors, one activation gate. Front Physiol. 2015;6:29. Epub 20150206. doi: 10.3389/fphys.2015.00029. PubMed PMID: 25705194; PubMed Central PMCID: PMCPMC4319557.

26. Doyle DA, Morais Cabral J, Pfuetzner RA, Kuo A, Gulbis JM, Cohen SL, et al. The structure of the potassium channel: molecular basis of K+ conduction and selectivity. Science. 1998;280(5360):69-77. doi: 10.1126/science.280.5360.69. PubMed PMID: 9525859.

27. Tao X, Hite RK, MacKinnon R. Cryo-EM structure of the open high-conductance Ca. Nature. 2017;541(7635):46-51. Epub 20161214. doi: 10.1038/nature20608. PubMed PMID: 27974795; PubMed Central PMCID: PMCPMC5500982.

28. Abuaku B, Boateng P, Peprah NY, Asamoah A, Duah-Quashie NO, Matrevi SA, et al. Therapeutic efficacy of dihydroartemisinin-piperaquine combination for the treatment of uncomplicated malaria in Ghana. Front Cell Infect Microbiol. 2022;12:1058660. Epub 20230106. doi: 10.3389/fcimb.2022.1058660. PubMed PMID: 36683700; PubMed Central PMCID: PMCPMC9853013.

29. Gibas KM, Kelly SG, Arribas JR, Cahn P, Orkin C, Daar ES, et al. Two-drug regimens for HIV treatment. Lancet HIV. 2022;9(12):e868–e83. Epub 20221026. doi: 10.1016/S2352-3018(22)00249-1. PubMed PMID: 36309038; PubMed Central PMCID: PMCPMC10015554.

30. Kerantzas CA, Jacobs WR. Origins of Combination Therapy for Tuberculosis: Lessons for Future Antimicrobial Development and Application. mBio. 2017;8(2). Epub 20170314. doi: 10.1128/mBio.01586-16. PubMed PMID: 28292983; PubMed Central PMCID: PMCPMC5350467.

31. Larkins-Ford J, Aldridge BB. Advances in the design of combination therapies for the treatment of tuberculosis. Expert Opin Drug Discov. 2023;18(1):83–97. Epub 20221228. doi: 10.1080/17460441.2023.2157811. PubMed PMID: 36538813; PubMed Central PMCID: PMCPMC9892364.

32. Okell LC, Drakeley CJ, Ghani AC, Bousema T, Sutherland CJ. Reduction of transmission from malaria patients by artemisinin combination therapies: a pooled analysis of six randomized trials. Malar J. 2008;7:125. Epub 20080709. doi: 10.1186/1475-2875-7-125. PubMed PMID: 18613962; PubMed Central PMCID: PMCPMC2491628.

33. Portsmouth S, Stebbing J, Gazzard B. Current treatment of HIV infection. Curr Top Med Chem. 2003;3(13):1458–66. doi: 10.2174/1568026033451808. PubMed PMID: 14529521.

34. Zahner H, Taubert A, Harder A, von Samson-Himmelstjerna G. Filaricidal efficacy of anthelmintically active cyclodepsipeptides. Int J Parasitol. 2001;31(13):1515–22. doi: 10.1016/s0020-7519(01)00263-6. PubMed PMID: 11595239.

35. Webb TI, Kshatri AS, Large RJ, Akande AM, Roy S, Sergeant GP, et al. Molecular mechanisms underlying the effect of the novel BK channel opener GoSlo: involvement of the S4/S5 linker and the S6 segment. Proc Natl Acad Sci U S A. 2015;112(7):2064–9. Epub 20150204. doi: 10.1073/pnas.1400555112. PubMed PMID: 25653338; PubMed Central PMCID: PMCPMC4343142.

36. Schewe M, Sun H, Mert Ü, Mackenzie A, Pike ACW, Schulz F, et al. A pharmacological master key mechanism that unlocks the selectivity filter gate in K. Science. 2019;363(6429):875-80. doi: 10.1126/science.aav0569. PubMed PMID: 30792303; PubMed Central PMCID: PMCPMC6982535.

37. Madeira F, Park YM, Lee J, Buso N, Gur T, Madhusoodanan N, et al. The EMBL-EBI search and sequence analysis tools APIs in 2019. Nucleic Acids Res. 2019;47(W1):W636–W41. doi: 10.1093/nar/gkz268. PubMed PMID: 30976793; PubMed Central PMCID: PMCPMC6602479.

38. Van Der Spoel D, Lindahl E, Hess B, Groenhof G, Mark AE, Berendsen HJ. GROMACS: fast, flexible, and free. J Comput Chem. 2005;26(16):1701-18. doi: 10.1002/jcc.20291. PubMed PMID: 16211538.

39. Lindorff-Larsen K, Piana S, Palmo K, Maragakis P, Klepeis JL, Dror RO, et al. Improved side-chain torsion potentials for the Amber ff99SB protein force field. Proteins. 2010;78(8):1950–8. doi: 10.1002/prot.22711. PubMed PMID: 20408171; PubMed Central PMCID: PMCPMC2970904.

40. Wang J, Wang W, Kollman P, Case D. Antechamber: an accessory software package for molecular calculations. J Am Chem Soc. 2001.

41. Wang J, Wolf RM, Caldwell JW, Kollman PA, Case DA. Development and testing of a general amber force field. J Comput Chem. 2004;25(9):1157–74. doi: 10.1002/jcc.20035. PubMed PMID: 15116359.

42. Friesner RA, Banks JL, Murphy RB, Halgren TA, Klicic JJ, Mainz DT, et al. Glide: a new approach for rapid, accurate docking and scoring. 1. Method and assessment of docking accuracy. J Med Chem. 2004;47(7):1739–49. doi: 10.1021/jm0306430. PubMed PMID: 15027865.

